# Characteristics of the C_4_ bundle sheath emerge in C_3_ rice after editing a plasma membrane proton ATPase

**DOI:** 10.64898/2026.05.22.727108

**Authors:** Lei Hua, Andrew R.G. Plackett, Na Wang, Julian M. Hibberd

## Abstract

C_4_ photosynthesis improves light, water and nitrogen-use efficiencies and can raise yield by 50% compared with the ancestral C_3_ pathway. Engineering C_4_ traits into C_3_ crops could substantially boost food production but requires coordinated modifications to leaf anatomy and cell-specific photosynthetic function. For example, C_4_ leaves contain more numerous, shorter bundle sheath cells that are photosynthetically active. In searching for transcriptional regulators of bundle sheath development in C_3_ rice, we unexpectedly found OSA3, a plasma membrane H^+^-ATPase that is expressed in bundle sheath cells as they elongate, and when knocked out reduces their length due to reduced apoplastic acidification. Bundle sheath cell number and chloroplast occupancy are increased. Thus, switching between C_3_ and C_4_ bundle sheath identity is controlled by acid growth, and OSA3 represents a simple tool for C_4_ engineering.

---

Photosynthesis supports life on Earth. Most plants use C_3_ photosynthesis, the fundamental biochemistry of which has remained unaltered since it evolved in bacteria (*1, 2*). In C_3_ species CO_2_ is fixed by Ribulose-1,5-Bisphosphate Carboxylase/Oxygenase (RuBisCO) in mesophyll cells. However, RuBisCO can also react with O_2_, initiating a series of repair reactions that waste energy and result in the loss of previously fixed carbon via photorespiration (*3, 4*). In response to reduced CO_2_ and water availability, multiple plant lineages have evolved carbon-concentrating mechanisms (*5*). One such example is the C_4_ pathway, where photosynthesis is partitioned between two compartments. Typically, in C_4_ plants, CO_2_ is first fixed in mesophyll cells into four carbon organic acids. After diffusion through plasmodesmata into bundle sheath cells decarboxylation of these acids releases high concentrations of CO_2_ around RuBisCO and suppresses photorespiration. Modifications to the biochemistry of photosynthesis in C_4_ leaves are accompanied by complex changes to leaf structure such as more veins, more plasmodesmata linking mesophyll and bundle sheath cells, more bundle sheath cells along the proximal-distal leaf axis, and also activation of chloroplast biogenesis in this tissue **(Fig. 1A and B)** (*5, 6*). These modifications to C_4_ leaves led to Hal Hatch, who discovered the pathway, to describe it as a unique blend of modified biochemistry, anatomy and ultrastructure (*7*). Despite the complexities of these changes to phenotype, C_4_ photosynthesis has evolved more than sixty times across the plant kingdom (*8*).

**Figure 1.**
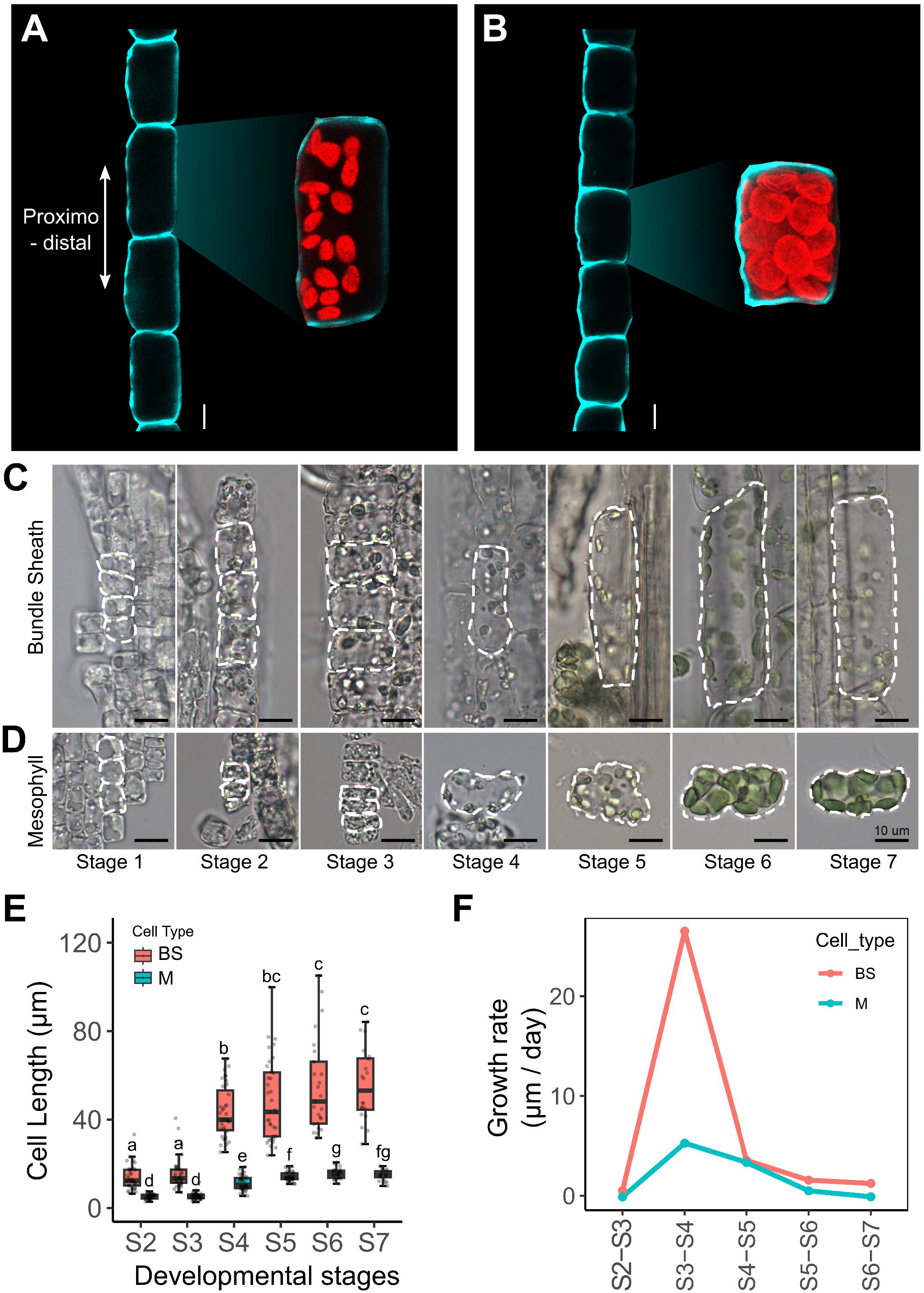
Rapid cell elongation along the proximo-distal leaf axis leads to C_3_ characteristics of the rice bundle sheath. (**A**,**B**) Key events associated with the evolution of C_4_ photosynthesis include greater numbers of less elongated bundle sheath cells with increased chloroplast occupancy. (**A**) C_3_ bundle sheath (*Oryza sativa*), (**B**) C_4_ bundle sheath (*Setaria viridis)*. Cell walls stained with Calcofluor White shown in turquoise, chloroplasts identified via chlorophyll autofluorescence in red. (**C**,**D**) Spatio-temporal gradient defining bundle sheath (**C**) and mesophyll development (**D**) in rice. Cells highlighted with white dashed lines. (**E**) Quantification of bundle sheath and mesophyll cell length along the proximo-distal leaf axis during development. (**F**) Growth rate of bundle sheath and mesophyll cells along the proximo-distal leaf axis during development. Scale bars: 10 μm in (**D**,**E**).

The extent to which the distinct phenotypes required for C_4_ to operate are underpinned by molecular changes that are genetically linked or unlinked has remained unclear. Some studies have predicted sequential alterations to phenotype, with each modification contributing to the pathway’s emergence by providing stable, fitness-neutral or advantageous traits (*5, 9, 10*). Other work proposes flexible and diverse evolutionary trajectories driven initially by non-photosynthetic drivers depending on ecological and developmental constraints (*11*). However, with the exception of the transcription factors *TOO MANY LATERALS1* that controls lateral vein production in leaves (*12*) and *GOLDEN2-LIKE* that orchestrates increased organelle and plasmodesmatal development in bundle sheath cells of C_3_ rice (*13*), there are few examples of genes known to control C_4_ traits. In particular, it remains unclear whether mutation of structural genes can lead to the simultaneous emergence of multiple C_4_-relevant traits. Identifying such a phenomenon has implications for attempts to engineering C_4_ photosynthesis into C_3_ crops to increase yield, water and nitrogen use efficiency (*14, 15*).

Previous work aiming to install the C_4_ pathway into rice has demonstrated production of C_4_ acids via Phospho*enol*pyruvate Carboxylase (*16, 17*), increased chloroplast occupancy of bundle sheath cells with greater plasmodesmatal linkages between mesophyll and bundle sheath cells (*13, 18*–*20*), and the production of more lateral veins (*12*). However, other phenotypes such as greater numbers of smaller bundle sheath cells along the proximo-distal leaf axis (*21, 22*) have not yet been engineered. We rationalised that a better understanding of bundle sheath development in C_3_ leaves would inform this process, and so first established a spatiotemporal developmental gradient for both bundle sheath and mesophyll cells. Through comparative analysis of gene expression and rates of cell development, we identified a structural gene with no known role in photosynthesis controlling the emergence of multiple traits characteristic of C_4_ photosynthesis. Specifically, loss-of-function alleles in a plasma membrane H^+^-ATPase had more bundle sheath cells that were shorter, and showed enhanced chloroplast occupancy. At the level of the leaf we estimate an ~30% additional bundle sheath cells are produced, and chloroplast number increased by ~21%. These findings provide new insights into the evolution of C_4_ photosynthesis and provide a simple roadmap to engineer bundle sheath cells in C_3_ crops to resemble those from C_4_ plants.

## Establishment of a spatiotemporal gradient of rice leaf development

To better understand bundle sheath cells in C_3_ species we identified and characterised a spatiotemporal gradient for their development in rice. By sampling the shoot apical meristem and leaf four as it develops, we identified key transitions associated with photosynthetic competency and specialisation. This revealed a two-step process comprising an initial period in which the blade grew but the sheath did not, and a second period where the sheath elongated (**Fig. S1A**). As leaf four was enclosed within the sheath of leaf three for nine days, dissection was used to sample it from day six onwards (**Fig. S1B**). We used this system to define a temporal developmental gradient for both bundle sheath and mesophyll cells, hereafter referred to as Stages 1-7 (**Fig. S1**). For example, at stage 1 bundle sheath and mesophyll cells were isodiametric, difficult to distinguish from one another and plastids were not easy to discern (**Fig. 1C and D**). Bundle sheath cells then became more elongated along the proximal-distal leaf axis, and by stages 2 and 5 respectively plastids became detectable and then green in both cell types (**Fig. 1C and D**). Chloroplast number per cell was not significantly different between bundle sheath and mesophyll cells (**Fig. S2D**), but as bundle sheath cells matured they showed the expected elongation and relatively low chloroplast occupancy characteristic of C_3_ plants (**Fig. 1C and E**). Mesophyll cells expanded much less along the proximal-distal leaf axis (**Fig. 1D and E**). At maturity, 75% and 25% of mesophyll and bundle sheath cell area respectively was occupied by chloroplasts (**Fig. S2G**). The difference between planar area occupied by chloroplasts in bundle sheath and mesophyll cells could be due to differences in cell growth, chloroplast biogenesis, or both. However, the number and size of chloroplasts was not statistically different between bundle sheath and mesophyll cells, and so total chloroplast area per cell was similar (**Fig. S2D-2F**). Because the area of bundle sheath cells was larger than that of the mesophyll, chloroplast occupancy in bundle sheath cells exhibited a negative correlation with cell area (**Fig. S3A**). This was not the case for mesophyll cells (**Fig. S3B**). The rate of increase in bundle sheath cell length and area was highest between stages 3 and 4 and was much greater than that of mesophyll cells (**Fig. 1F, Fig. S4**).

To provide insight into the molecular basis of these phenotypes we undertook laser capture microdissection to allow deep sequencing of both cell types (*23*). Although these cell types were difficult to distinguish at early stages, starch in bundle sheath cells (*24*) allowed them to be identified, dissected and RNA isolated (**Fig. 2A and B, Fig. S5A, B, D-F, Fig. S6**). Whilst genes associated with cellular respiration and solute transport were upregulated in the bundle sheath, those relating to protein biosynthesis, coenzyme biosynthesis and secondary metabolism were more highly expressed in mesophyll cells (**Fig. S7**). These patterns of gene expression align with the known function of the bundle sheath in solute transport and the mesophyll in photosynthesis (*25*), strongly suggesting differentiation had taken place by stage 2.

**Figure 2.**
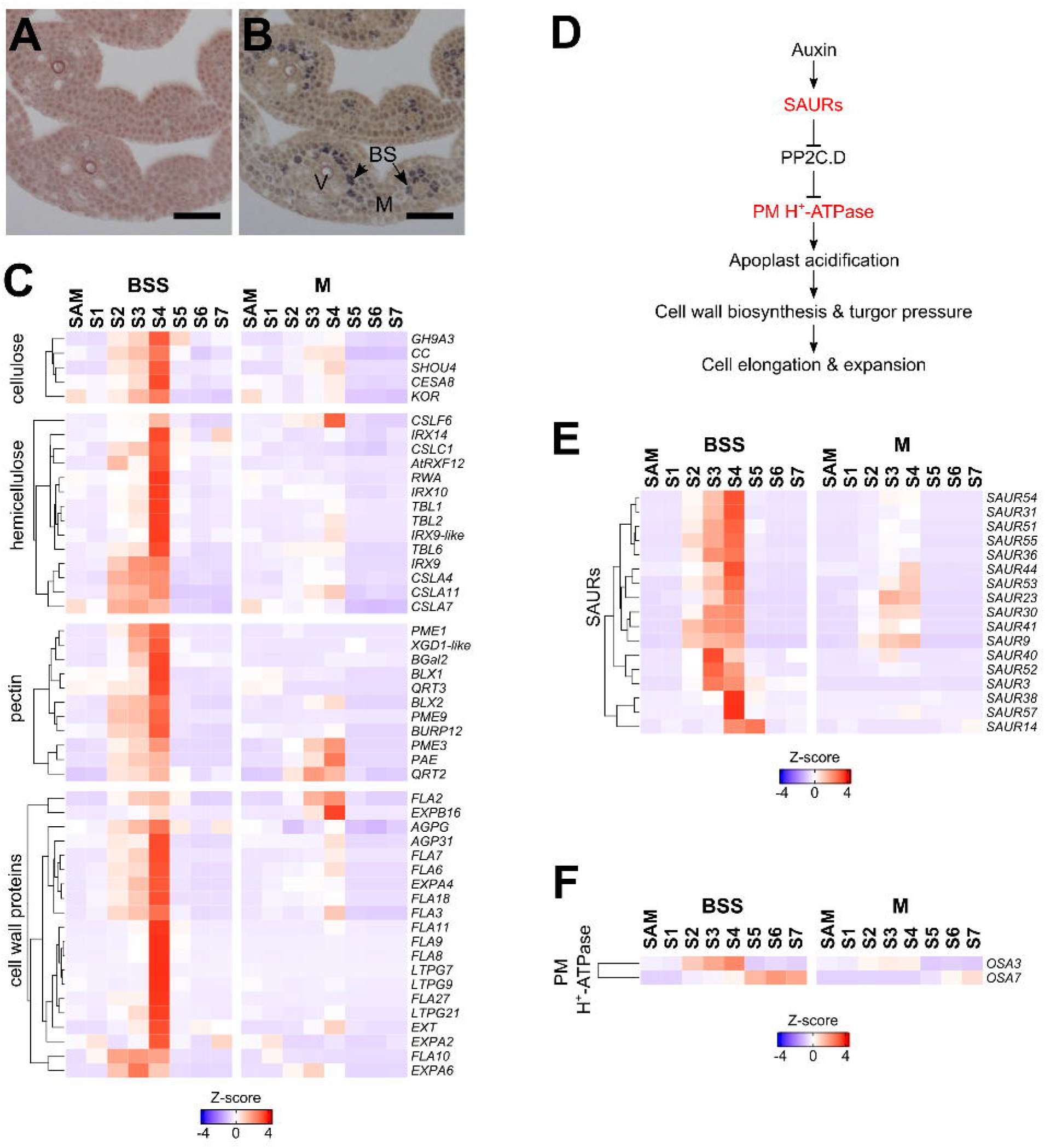
Rapid bundle sheath elongation is correlated with transcripts involved in apoplast acidification. (**A**,**B**) Starch staining allowed bundle sheath cells to be distinguished from mesophyll cells at early developmental stages. Cross sections at stage 2 before (**A**) and after (**B**) starch staining. Scale bars represent 50 μm. (**C**) Expression of cell wall biogenesis-related genes from modules 14 and 9. (**D**) Schematic summarising auxin-mediated apoplast acidification. (**E**,**F**) Expression of transcripts encoding Small Auxin Upregulated RNAs (SAURs) (**E**), and Plasma Membrane (PM) H^+^-ATPases *OSA3* and *OSA7* (**F**).

Weighted Gene Co-expression Network Analysis (WGCNA) identified twenty-two unique co-expression modules that either showed temporal or spatial peaks in expression (**Fig. S8**). Notably, substantial cell-specific gene expression had already been established at early developmental stages. Module 15 and 17 consistently exhibited expression in bundle sheath strands as early as stage 2 **(Fig. S8)**. Similarly, all modules preferentially expressed in mesophyll cells (M18–M22) were active during stages 2 to 4 (**Fig. S8)**. We linked these spatiotemporal patterns in gene expression to biological functions by assessing the extent to which Mercator categories were enriched in each module. Modules 1 to 5 contained genes most highly expressed in the SAM and leaf primordia and were significantly enriched in RNA biosynthesis, RNA processing and chromatin organisation (**Fig. S8, S9B**). Module 6 was associated with processes related to cell division, chromatin organisation, cytoskeleton organisation and DNA damage response and expressed in both cell types up to stage 4 (**Fig. S8, S9B**). Consistent with the early onset of photosynthesis during leaf development in mesophyll cells(*26*) photosynthesis-related genes were highly overrepresented in modules 19-22 and preferentially expressed in mesophyll in either early and/or late stages (**Fig. S8, S9B**). Similarly, bundle sheath strands were associated with modules 16 and 17 that were enriched in solute transport and phytohormone action (**Fig. S8, S9B**). The spatiotemporal gene expression patterns obtained thus successfully correlated strongly with observed morphologies and functions of the two cell types.

### Rapid bundle sheath elongation is correlated with patterning of genes associated with apoplast acidification

To better understand how structure and function of mature bundle sheath cells is established, we focussed on the five modules that were positively and highly correlated with their changes in cell length and area **(Fig. S10A)**. Modules 14 and 9 were most strongly enriched in genes associated with cell wall organisation (**Fig. S9B, S10B**) including transcripts encoding cellulase, hemicellulose, pectin biosynthetic enzymes and cell wall proteins (**Fig. 2C**). Consistent with previous analysis of hypocotyl epidermal cells (*27*–*29*) genes associated with primary active transport including P-type and V-type H^+^-ATPases, plasma membrane intrinsic proteins (aquaporins), and auxin signalling were also present in modules associated with rapid bundle sheath cell elongation (**Fig. S10B**). Auxin promotes cell expansion by promoting transcription of Small Auxin Up RNA (SAUR) genes, which then activate plasma membrane H^+^-ATPases by inhibiting PP2C.D phosphatase activity (*27*) (**Fig. 2D**). The SAUR family was statistically significantly over-represented in module 9 (Chi square test *P* < 0.0001) and module 14 (Chi square test *P* < 0.0001). Remarkably, seventeen SAURs in these modules showing progressively increased expression in the bundle sheath strands from stage 2 to stage 4 (**Fig. 2E, Fig. S11A and B**). Moreover, two plasma membrane H^+^-ATPases showed strong and specific expression in bundle sheath cells. *OSA3* (LOC_Os12g44150) was increasingly expressed in the bundle sheath strands from stage 2 to stage 4 (**Fig. 2F, Fig. S11C**), while *OSA7* (LOC_Os04g56160) peaked in expression from stage 5 onwards (**Fig. 2F, Fig. S11C**).

Consistent with the pattern of transcripts associated with apoplastic acidification, leaf tissue from stages 2 to 5 led to acidification of media (**Fig. S12A and B**), and this was increased or reduced after application of pharmacological agents known to activate or block activity of plasma membrane H^+^-ATPases (**Fig. 3A and B**). Length but not width of bundle sheath cells also increased or reduced after application of these agents (**Fig. 3C and D, Fig. S12D**). While application of the auxin polar transport inhibitor N-1-naphthylphthalamic acid had no detectable impact on bundle sheath cell size (**Fig. S12E**), synthetic auxin increased length of bundle sheath cells (**Fig. S12E**) and inhibitors of auxin biosynthesis resulted in shorter cells (**Fig. S12E**). These results suggest that activity of plasma membrane localised H^+^-ATPases control elongation of bundle sheath cells in response to local auxin biosynthesis, rather than auxin transport.

**Figure 3.**
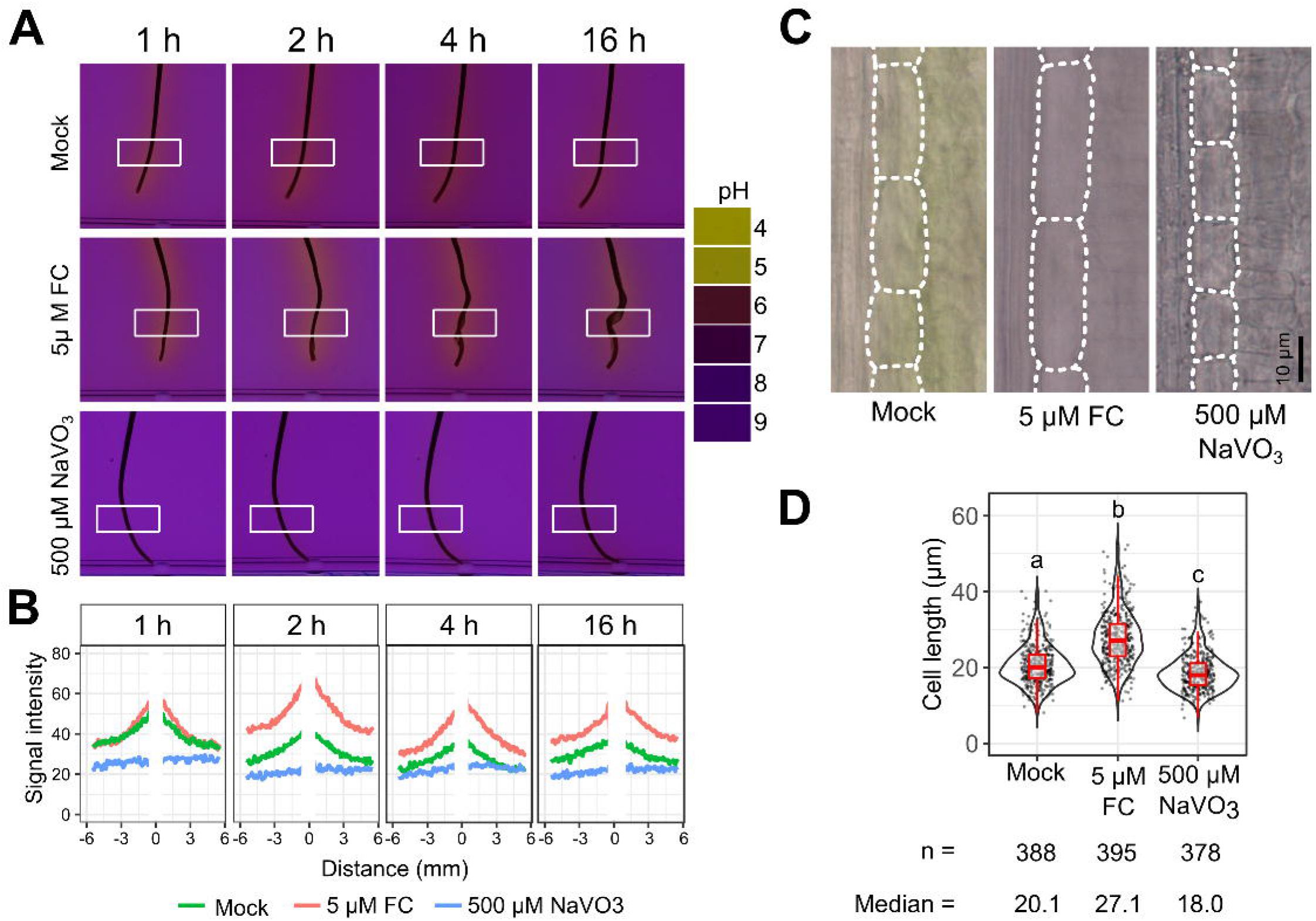
Plasma membrane H^+^-ATPase regulates bundle sheath elongation. (**A**) Fusicoccin (FC) enhances, and sodium vanadate (NaVO_3_) suppresses acidification around stage 3 rice leaves. (**B**) Quantification of acidification after FC and NaVO_3_ treatment. (**C**) Representative images of bundle sheath cells along the proximo-distal leaf axis after treatment with FC or NaVO_3_. Bundle sheath cells indicated with dashed lines, scale bars represent 10 μm. (**D**) Quantification of bundle sheath cell length after each treatment.

### Characteristics of the C_4_ bundle sheath emerge after editing *OSA3*

As expression analysis and pharmacological treatments implicated apoplastic acidification in rapid bundle sheath cell elongation we tested this genetically. Loss-of-function alleles for the *OSA3* and *OSA7* genes were generated by gene editing (**Fig. S13**). *osa3* and *osa7* mutants contained shorter bundle sheath cells (**Fig. 4A and B** and **Fig. S14A and B**). Leaves of *osa7* were shorter than controls and plants stunted (**Fig. S14I, Fig. S15C, D and F)** but this was not the case for *osa3* (**Fig. S15A, B &E, Fig. S16A**), and so we prioritised analysis of the latter. Width of bundle sheath cells in *osa3* mutants slightly increased (**Fig. S16A**) but due to the reduction in length, overall, bundle sheath cell planar area was decreased (**Fig. S16B**). *osa3* mutants exhibited reduced sensitivity to fusicoccin that activates H^+^-ATPases (**Fig. 4C, Fig. S16C and D**) consistent with bundle sheath length being determined by plasma membrane H^+^-ATPase activity. Although total planar chloroplast area in bundle sheath cells was similar in *osa3* and wild type (**Fig. S17B-E**), the reduction to cell area meant that chloroplast coverage per cell was increased (**Fig. 4D and E**). Vein number was unaffected in *osa3* mutants (**Fig. S17F**) and so we calculate that *osa3* leaves contain ~30% more bundle sheath cells than wild type (**Fig. S17H**). The unchanged planar chloroplast area per cell and the greater number of bundle sheath cells along each vein means that *osa3* mutants contained ~21% more bundle sheath chloroplasts (**Fig. S17I**) and ~34% greater bundle sheath chloroplast area than controls **(Fig. 4F)**.

**Figure 4.**
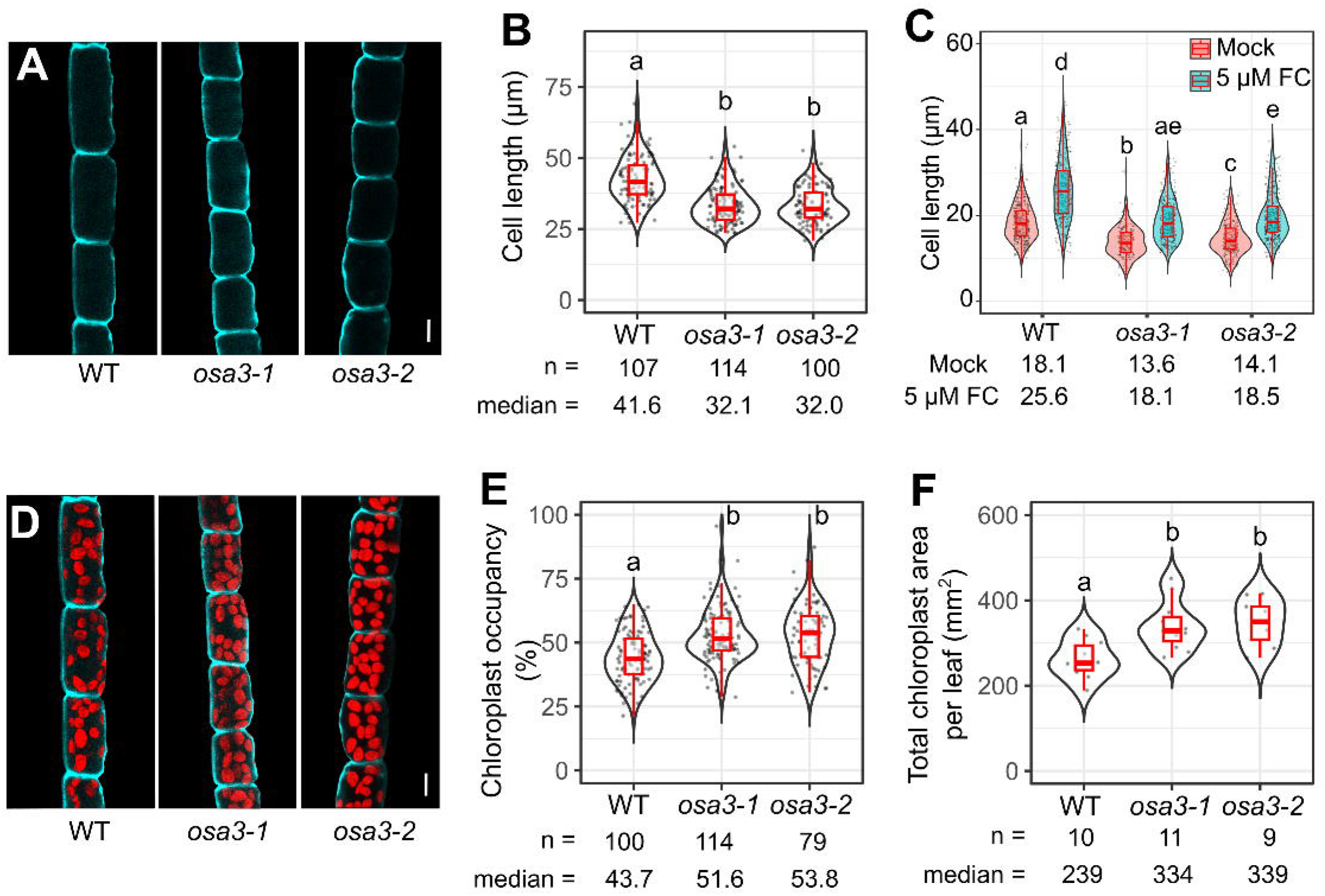
Editing plasma membrane H^+^-ATPases generates characteristics of C_4_ bundle sheath cells. (**A**) Representative images of bundle sheath cells in wild type and *osa3* mutants, cell walls visualized in turquoise using Calcofluor White staining. (**B**) Quantification of bundle sheath length in wild type and *osa3* mutants. (**C**) the bundle sheath in *osa3* mutants at stage 3 exhibits reduced sensitivity to FC. (**D**) bundle sheath cells with chloroplasts shown by chlorophyll fluorescence in red. (**E**) Quantification of chloroplast occupancy in wild type and *osa3* mutants. (**F**) total chloroplast area in bundle sheath cells of leaf 4 blade. Scale bars represent 10 μm in **A** and **D**.

Overall, our data support a model in which elongation of bundle sheath cells in rice is regulated by plasma membrane associated H^+^-ATPases. Loss-of-function alleles for *OSA3* contain more and shorter bundle sheath cells along the proximo-distal leaf axis, two characteristics of C_4_ plants. Additional emergent properties of this simple genotypic change include increased chloroplast occupancy per bundle sheath cell and more bundle sheath chloroplasts in the leaf. Thus, the single *OSA3* gene controls multiple properties associated with the evolution of C_4_ photosynthesis, and loss-of-function alleles provide a chassis into which additional C_4_ components could be engineered.

## Discussion

Our analysis of rice bundle sheath cells provides new insight into the evolution of C_4_ photosynthesis, attempts to engineer C_4_ photosynthesis into C_3_ crops to increase yield, and also mechanisms controlling elongation of a specific cell type embedded deep inside leaf tissue, Constrained by the cell wall, elongation of plant cells is regulated by multiple factors, including turgor pressure, cell wall extensibility, and the synthesis and integration of new wall components (*30, 31*). In the early 1970s analysis of sunflower hypocotyls led to the acid growth hypothesis (*32, 33*) that posited auxin-induced apoplastic acidification promotes cell wall loosening and elongation. Pharmacological studies, particularly with fusicoccin, reinforced this model by demonstrating that activation of plasma membrane H^+^-ATPases can trigger proton efflux and drive cell expansion (*34*). More recently, genetic analysis revealed that auxin perception promotes transcription of *SAUR* genes, which both activate plasma membrane H^+^-ATPases through phosphorylation, and inhibit PP2C.D phosphatases that repress H^+^-ATPase activity. This auxin-dependent, SAUR-mediated activation of plasma membrane H^+^-ATPases leads to rapid cell expansion (*27*–*29*). However, the identity of the cell types responding to acid growth remained unclear. Observations are largely limited to surface tissues, such as the epidermis of elongating hypocotyls, epicotyls and roots, where acidification and growth responses can be readily measured (*31*), and so how internal tissues respond to apoplastic acidification is less well understood. Moreover, to our knowledge, genetic evidence for acid growth in leaf tissues, particularly in monocotyledons such as rice, is lacking. While mutation of *OSA7* which is preferentially expressed in bundle sheath cells during later stage of maturation led to shorter bundle sheath cells, additional pleiotropic effects on growth were detected. As plasma membrane H^+^-ATPases also contribute to a wide range of physiological processes, including nutrient uptake, stomatal opening, phloem loading, and plant-pathogen interactions (*35*–*37*), effects beyond the control of anisotropic bundle sheath cell expansion is not unexpected. However, in the case of *OSA3*, which is preferentially expressed during early bundle sheath maturation, the effect on elongation of these cells did not have detectable impacts on whole plant growth. Our pharmacological and genetic analysis implicate *OSA3* specifically in bundle sheath elongation, and gene expression in the bundle sheath implies that the other components associated with acid growth and cell wall relaxation and synthesis act in concert with this particular plasma membrane H^+^-ATPase. To our knowledge, these findings provide the first direct evidence that acid growth contributes to shaping internal leaf structure in a monocotylenous species.

Our findings now also implicate plasma membrane H^+^-ATPases in the evolution of C_4_ photosynthesis. These proteins have been linked to processes related to photosynthesis such as hydraulic conductance and stomatal opening (*38*–*41*). Here we report that multiple characteristics associated with evolution of the C_4_ pathway emerge after generating loss-of-function alleles for the *OSA3* gene encoding a bundle sheath preferentially expressed plasma membrane ATPase in C_3_ rice. Light activates plasma membrane H^+^-ATPases (*38*–*41*), and delayed exposure to light leads to shorter bundle sheath cells (*42*), consistent with a key role for plasma membrane H^+^-ATPases in controlling their size. Of specific relevance to C_4_ evolution, the data indicate that deleterious mutations to the function of this protein caused shorter bundle sheath cells along the proximo-distal axis, but leaf length remained unaffected. The outcome is that the number of bundle sheath cells is increased significantly compared with wild type - a key feature needed for C_4_ photosynthesis. Perturbing bundle sheath cell expansion therefore led to a compensatory increase in cell number, and feedback between cell expansion and division. Crosstalk between cell division and expansion is well documented in epidermal layers, where constraining cell expansion can increase cell division to maintain total organ size (*43*–*45*). Our findings support the notion that this mechanism also operates in specific internal tissue layers of the leaf.

Coordination between cell and chloroplast division (*46, 47*) means that chloroplast number per bundle sheath cell is not impacted by loss of *OSA3* function. Moreover, the reduced elongation of each bundle sheath cell in *osa3* mutants leads to their planar area being smaller, and so the proportional area of each cell occupied by chloroplast is increased. These linked phenomena mean that *osa3* rice leaves contain a significant increase in bundle sheath chloroplast area, and higher ratio of bundle sheath to mesophyll chloroplast compared with wild type. Activation of photosynthesis in bundle sheath cells is a fundamental characteristic needed for C_4_ photosynthesis (*5, 48, 49*). We conclude that mutation to plasma membrane localised proton ATPases could have facilitated the evolution of C_4_ traits in lineages that have evolved the C_4_ pathway.

In addition to providing insights into how C_4_ traits may have evolved, discovering that *OSA*3 controls bundle sheath elongation and chloroplast occupancy has implications for efforts to improve crop productivity and nitrogen and water use through C_4_ engineering. For example, *osa3* that contains more bundle sheath cells along veins could be combined with the *tml1* mutant that contains up to 23% more bundle sheath cells around veins (*12*). *osa3* and *tml1* should therefore increase bundle sheath cell number by up to 53%. Moreover, we anticipate that the 21% increase in chloroplast occupancy per bundle sheath cell of *osa3*, could be further enhanced by combining with lines over-expressing *GOLDEN2-LIKE* in which direct manipulation of chloroplast biogenesis increased individual chloroplast area per cell (*13*). Collectively, we anticipate that these three genetic perturbations provide a useful chassis in which the rice bundle sheath is activated for photosynthesis. This could then be used to accept biochemical components that allow the C4 cycle to operate.

In summary, our spatiotemporal analysis of early bundle sheath development identifies apoplast acidification mediated by the plasma membrane H^+^-ATPase OSA3 as critical for cell elongation. Analysis of loss-of-function osa3 lines identify a gene that could have been co-opted during evolution of initial C4 traits prior to emergence of the full C4 pathway, and also establishes a promising framework from which to engineer C4 photosynthetic into C3 crops.

## Material and methods

### Plant growth conditions, laser capture microdissection and RNA sequencing

Kitaake (*O. sativa* ssp. *japonica*) seeds were germinated and grown in mixture of 1:1 topsoil and sand in a growth cabinet with 12h day/12h night photoperiod, 28 °C day/25 °C night temperature, relative humidity of 60%, and photon flux density of 400 μmol m^−2^ s^−1^. Plants were supplemented with Everris Peters Excel Cal-Mag Grower N.P.K. 15-5-15 fertiliser (LBS Horticulture, UK) and DTPA chelated iron (Gardening Direct, UK).

For laser capture microdissection, leaf blade from 5-mm above leaf ligule at respective developmental stages was separated from midrib and then embedded in Steedman’s wax as previously describe (*23*). Microtome was used to create 7 μm paradermal sections which were then mounted on Polyethylene Naphthalate (PEN) membrane slides using DEPC treated water (*23*). Steedman’s wax was removed by exposing the slides to 100% acetone for 1 minute prior to LCM, to distinguish bundle sheath in younger leaf tissue (stage 2, 3, and 4), slides were incubated in 1% iodine in acetone for 1 min. Using the Arcturus LCM platform, isolated cells were collected on CapSure™ Macro Caps, and RNA was extracted using the PicoPure RNA Isolation Kit with on-column DNase I treatment. RNA quality and concentration were analysed using an Agilent Bioanalyser RNA 6000 Pico assay. RNA from at least three different leaves were combined into one biological replicate, 1.4-3ng of bundle sheath strands or mesophyll RNA with three to four biological replicates for each stage were reverse transcribed and amplified using the SMART-Seq v4 Ultra Low Input RNA Kit (Clontech) according to the manufacturer’s instructions, 11 PCR cycles were used for all samples. 1 ng of amplified cDNA was used as input for preparing library using the Nextera XT DNA Library Preparation Kit (Illumina). Libraries were sequenced using Illumnia’s NovaSeq 6000 sequencing platform to generate 31-50 million paired-ended 150-bp reads for each sample.

### Differential gene expression and co-expression network analysis

The leading 15-bp and tailing 3-bp of raw reads were trimmed and reads with a quality score of <10 and shorter than 20-bp as well as adaptor sequences were removed using ‘bbduk.sh’ script from BBMap 38.91 (*46*). Transcript abundance was determined after the remaining reads were quantified using salmon 1.5.1 (*47*) against *Oryza sativa* v7.0 transcripts from Phytozome V13 (*48, 49*) with selective alignment (“--validateMappings”) enabled. Gene-level abundance (transcripts per million, TPM) was summarized using tximport 1.18.0 (*50*), genes with TPM > 1 in at least three samples of at least one developmental stage were defined as expressed genes, resulting in 22753 genes. To evaluate transcriptome differences across samples using these genes, a principal component analysis (PCA) analysis was performed using the ‘prcomp’ function. Using the Mapman categories annotated by Mercator4 (*51*), the top and bottom 5% of loading genes of the first three principal components were analysed for over-representation via Fisher’s exact test with FDR threshold of 0.1. Gene-level counts were used for differential gene expression analysis using edgeR (v 3.40.2) (*52*), false discovery rate (FDR) of <0.05 were used to define differentially expressed genes. Log_2_ transformed TPM from expressed genes were used for co-expression network analysis using weighted correlation network analysis (WGCNA (v.1.63)) (*53*) with soft threshold of 12, minimal module size of 100 genes, MEDissThres cut-off of 0.1. To quantify module and trait associations, average cellular and chloroplastic growth rates were calculated as the changes of median values per day between adjacent stages, Pearson’s corelation between growth rates and module eigengenes was performed according to Tutorials for the WGCNA package (https://horvath.genetics.ucla.edu/html/CoexpressionNetwork/Rpackages/WGCNA/Tutorials/) using a cut-off of *P* value of 0.1.

### In vitro gel system for apoplast acidification, and pharmacological interventions

The solution containing 10 mM KCl, 1 mM CaCl_2_, and 90 mg/L bromocresol purple (*54, 55*) was adjusted for pH at 7.0 using 1 M KOH, Ultrapure LMP (low melting point) agarose (Invitrogen) was added to 0.5% (w/v) and microwaved to dissolve. Activator (*56, 57*) or inhibitor (*58, 59*) of plasma membrane H^+^-ATPase were added to the medium when it was cooled down to around 37 °C and the pH was re-adjusted to 7.0 using 100 mM KOH. The medium was poured into Petri dishes with 5 mm depth and cooled down to room temperature. Young fourth leaves at different stages were carefully dissected and gently pressed into semi-solidified agarose gel. Agarose gel with leaves were placed in Panasonic light incubator with light setting of 3LS at 28 °C. Plates were photographed with a Cannon EOS 750D digital camera on a light box and photos were processed with ImageJ (version 1.52a) (*60*). To quantify the effect of yellowing around leaf tissue caused by acidification, images were split into channels. Using the green channel image, a 2 × 10 mm region were selected at the base of leaf using the rectangle tool. The “Analyze > Plot profile” tool was used to obtain column average of grey values.

For pharmacologial treatments, Kitaake seeds were germinated and grown on ½ MS agar media in a 9-cm magenta box for ten days to reach stage 3 in a Panasonic growth chamber at 28 °C and in 12h day/12h night photoperiod. Test reagents were added to ½ MS basal salt solution with 0.1% Tween-20 and 50 mg/L Timentin, the pH was adjusted to 7.0 with 0.1 M KOH. Seedlings were removed from growing media without damaging the root, coleoptile and the first two leaves were carefully removed with a sharp tweezer, then the fourth leaves at stage 3 (10-14 cm in length) were slowly pulled out of leaf 3 sheath until ~ 5 mm of its base, then the fourth leaves with the third leaves attached were placed into 50 ml Falcon tubes containing 30 ml ½ MS solution with test reagents, this ensured that ~5-cm leaf base were bathed in solution, top of falcon tubes were sealed with parafilm with the third leaf outside. Each tube with ten seedlings were placed in Panasonic light incubator with weak light (3LS) at 28 °C for 48 hours. Bundle sheath and mesophyll cells at 5-mm above ligule of the fourth leaves were imaged under an Olympus BX41 light microscope with a mounted Micropublisher 3.3 RTV camera (Q Imaging), photos were captured with Q-Capture Pro 7 software.

### Generation of CRISPR knockout mutants and quantification of cell dimensions and chloroplast content

To generate the CRISPR constructs, two guide RNA for each gene targeting the second or third exon of OSA3 and OSA7 were designed using the online guide RNA selection tool CRISPOR (https://crispor.gi.ucsc.edu/). Two guide RNA targeting *OSA3* (5’-CGAGGGTCTCACAACTCAGC-3’, 5’-AGCTATCATGGCCATTGCGC-3’), two guide RNA targeting *OSA7* (5’-CAGCTGAAATGTACGCGCGA-3’, 5’-GGCCGAACATCTCGATACGC-3’) were cloned into golden gate level 1 vector in which *OsU3* promoter driving above guide RNA and scaffold at the R3 or R4 position. Then they are assembled into level 2 vector with *ZmUBI* promoter driving Cas9p. The resulting level 2 vectors were transformed into *Agrobacterium tumefaciens LBA4404*.

Rice transformation was performed as described previously (Hiei et al., 2008) with the following modifications. Kitaake seeds were dehulled and sterilised with 2.5% (v/v) sodium hypochlorite for 15 mins, calli were induced on NB medium with 2 mg/L 2,4-D at 30 °C in darkness for 3-4 weeks. Actively growing calli were then co-incubated with *A. tumefaciens* described above in darkness at 25 °C for 3 days, then selected on NB medium supplied with 35 mg/L hygromycin B for 4 weeks. Proliferating calli were subsequently transferred to NB regeneration medium containing 2 mg/L kinetin, 0.2 mg/L 1-naphthaleneacetic acid and 0.8 mg/L 6-benzylaminopurine for four weeks at 28 °C under constant light. Plantlets were transferred to NB rooting medium containing 0.1 mg/L 1-naphthaleneacetic acid and incubated in Magenta pots for two weeks at 28 °C in the light. Finally, plants were transferred to a 1:1 mixture of topsoil and sand and grown in a growth cabinet in 12 h light/12 h dark photoperiod, at temperature of 28 °C at day 20 °C at night, with a relative humidity of 65% and a photon flux density of 400 μmol m^−2^ s^−1^. Editing of target genes were confirmed using sanger sequencing at T0 generation, Cas9-free homozygous edited plants were selected at T1 generation.

To quantify changes in cell dimension and chloroplast across the developmental gradient, leaf tissue at respective stages was processed as described (*38*). In brief, leaf tissue was harvested and fixed with 4% wt/vol paraformaldehyde in 1x phosphate buffered saline (PBS) buffer (pH 6.9) overnight, then fixative were replaced with 1x PBS solution (pH 6.9) and stored at 4°C. Cell walls were digested using 0.2 M sodium-EDTA solution (pH 9.0) at 55°C for 2 h followed by 2% wt/vol Aspegillus niger pectinase at 45°C for 2 h. Individual cells were imaged within 24 h of cell wall digestion using an Olympus BX41 microscope and an Olympus PLN ×20 objective lens, recording each cell at 4-5 separate focal depths to accurately measure the morphology of individual chloroplasts throughout the cell volume. Images were captured using MicroPublisher camera and QCapture Pro 7 software (Teledyne Photometrics). Cell and chloroplast measurements were taken from scaled images using FIJI (Schindelin et al., 2012).

To compare cell dimensions and chloroplast content between CRISPR knockout mutants and the null segregants, middle section (2-cm) of fully expanded fourth leaves were dissected and infiltrated with 1% (w/v) glutaraldehyde in 1X PBS buffer. Samples were left in that solution for overnight and then washed with 1X PBS buffer. Before imaging, the adaxial side of fixed leaf was scrapped gently with razor blade as described (*61*). The scrapped leaf fragment was stained with calcofluor white (0.1%; Sigma) for 1 min and then rinsed with H2O. Confocal imaging was performed using a Leica SP8X confocal microscope upright system (Leica Microsystems) under a 40x water immersion objective. Calcofluor white was excited at 405 nm and emitted fluorescence captured from 452-472 nm. Chlorophyll autofluorescence was excited at 488 nm and emission captured 672–692 nm. Z-stacking images of the bundle sheath were taken from three intermediate veins of each leaf sample, twelve to twenty individual plants per mutant were analysed. Maximum intensity projection of Z-stack images was generated in ImageJ (v1.54g) which was used to quantify bundle sheath cell dimensions, chloroplast size, number and total chloroplast area in each bundle sheath cell. Around ten bundle sheath cells per plant were analysed for bundle sheath dimensions, and around ten bundle sheath cell per plant and five to ten plants in total were analysed for chloroplast content.

## Statistics and reproducibility

For the statistical analysis of leaf parameters, data normality was assessed using the Shapiro-Wilk test. If the data were not normally distributed across all samples, pairwise Wilcoxon rank-sum tests were conducted using the pairwise.wilcox.test function from the stats package (v4.2.0) in R v4.2.3, with *p*-value adjusted by the Benjamini-Hochberg correction, A false discovery rate (FDR) threshold of < 0.05 was used to determine statistical significance. Data were visualized as violin plots, the 25^th^, 50^th^ and 75^th^ percentile was shown in boxplot with whiskers of 1.5x interquartile range, plots were generated using ggplot2 (v3.5.0).

## Author contributions Statement

L.H. and J.M.H. conceived the work. J.M.H. guided execution of experiments and oversaw the project. A.R.G.P established the spatiotemporal gradient of rice leaf and collected samples; L.H. performed LCM, RNA-seq data analysis and characterisation of mutants, N.W. conducted rice transformation. L.H. and J.M.H. wrote the manuscript with input from all authors.

## Data, code, and materials availability

Code associated with this manuscript is available in the Github repository: https://github.com/hibberd-lab/. RNA sequencing data are available under the BioProject accessions PRJNA1205924.

## Supplementary Materials

Materials and Methods

Figs. S1 to S17

Data S1 to S6

