## Supplemental Data for "Characteristics of the C_4_ bundle sheath emerge in C_3_ rice after editing a plasma membrane proton ATPase"

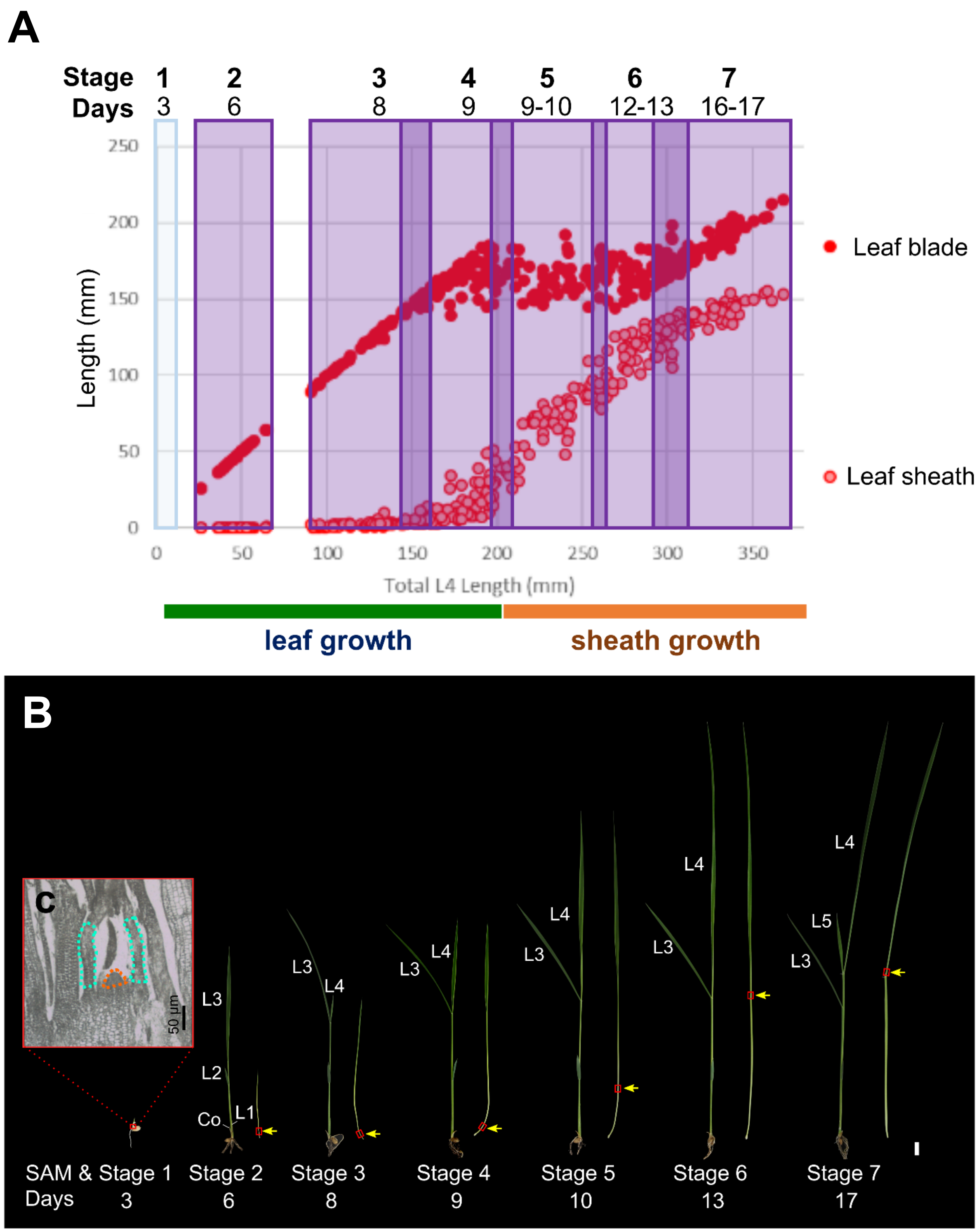


**Supplementary Figure 1. Establishing a spatiotemporal gradient for bundle sheath and mesophyll cell development in rice.**
(**A**) Growth of the blade and sheath of the fourth leaf during maturation. (**B**) Representative images showing staging of plants sampled to establish the gradient. For each stage, the fourth leaf (right side of each plant) was dissected. Yellow arrows indicate ligules of the fourth leaf, red boxes represent regions of sampling. Co represents coleoptile, L1 to L4 represent the first to the fourth leaves. (**C**) Section showing the base of a 3-day-old seedling, with the shoot apical meristem (SAM) and leaf primordia (Stage 1) highlighted with orange or turquoise dashed lines. Scale bars represent 5-mm in **B**, 50 µm in **C**.

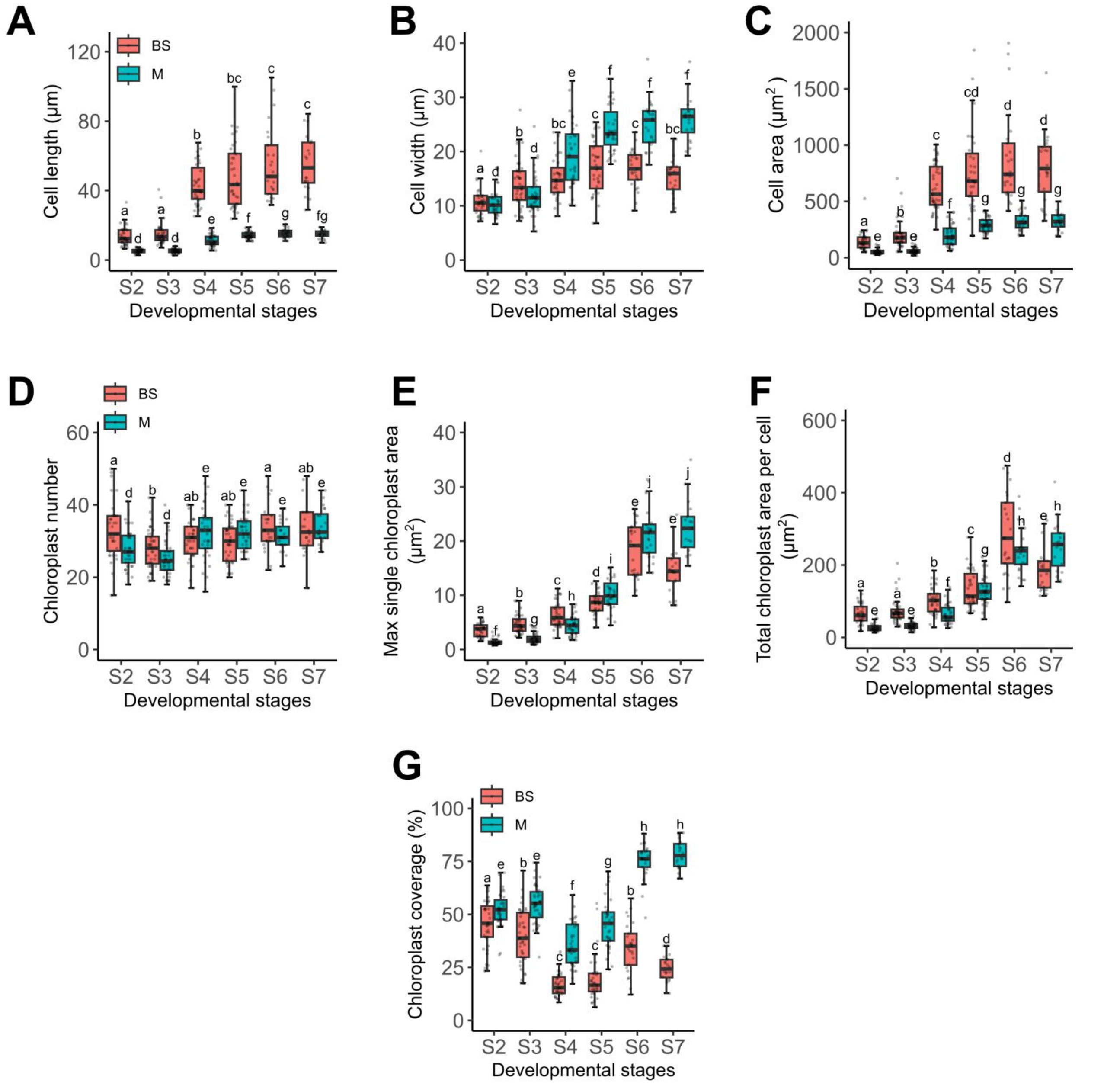


**Supplementary Figure 2. Quantification of parameters associated with bundle sheath and mesophyll cell development**.

Quantification of cell length along the proximodistal axis (**A**), cell width (mediolateral axis) (**B**), cell area (**C**), chloroplast number (**D**), maximum area of individual chloroplasts (**E**), total planar chloroplast area per cell (**F**), and chloroplast coverage per cell (**G**).

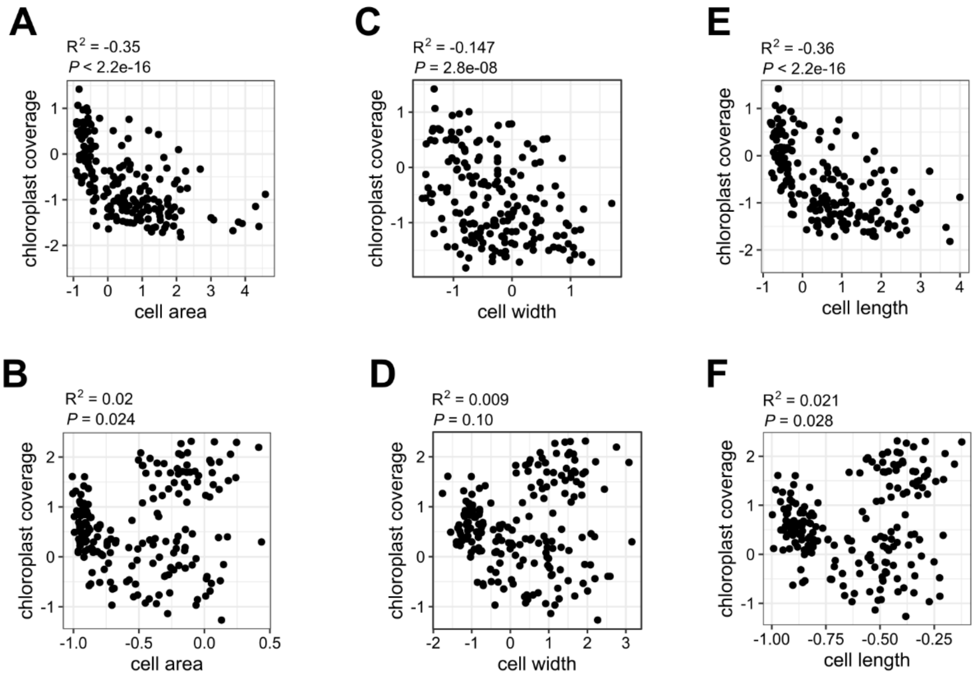


**Supplementary Figure 3. Relationships between cell dimensions and chloroplast coverage of each cell type.**

Correlation analysis of cell area (**A**,**B**), cell width (**C**,**D**) and cell length (**E**,**F**) for the bundle sheath (**A**,**C**,**E**) and mesophyll (**B**,**D**,**F**). Data were normalised using Z-scores.

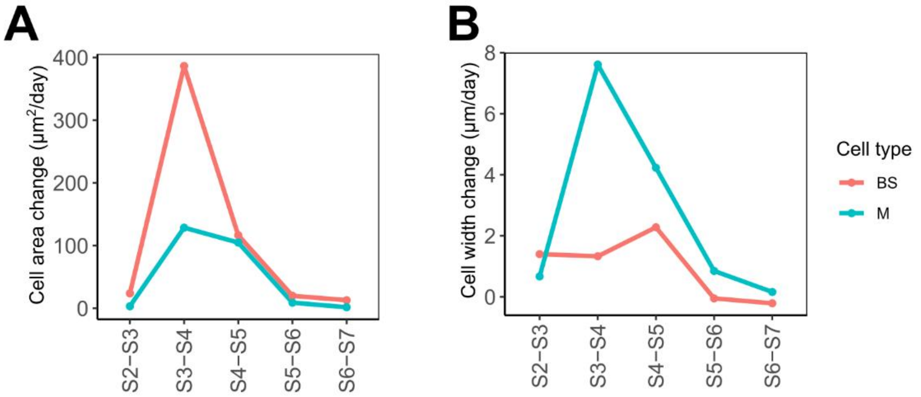


**Supplementary Figure 4. Rate of growth measured in cell area (A) and cell width (B) during leaf four maturation.**

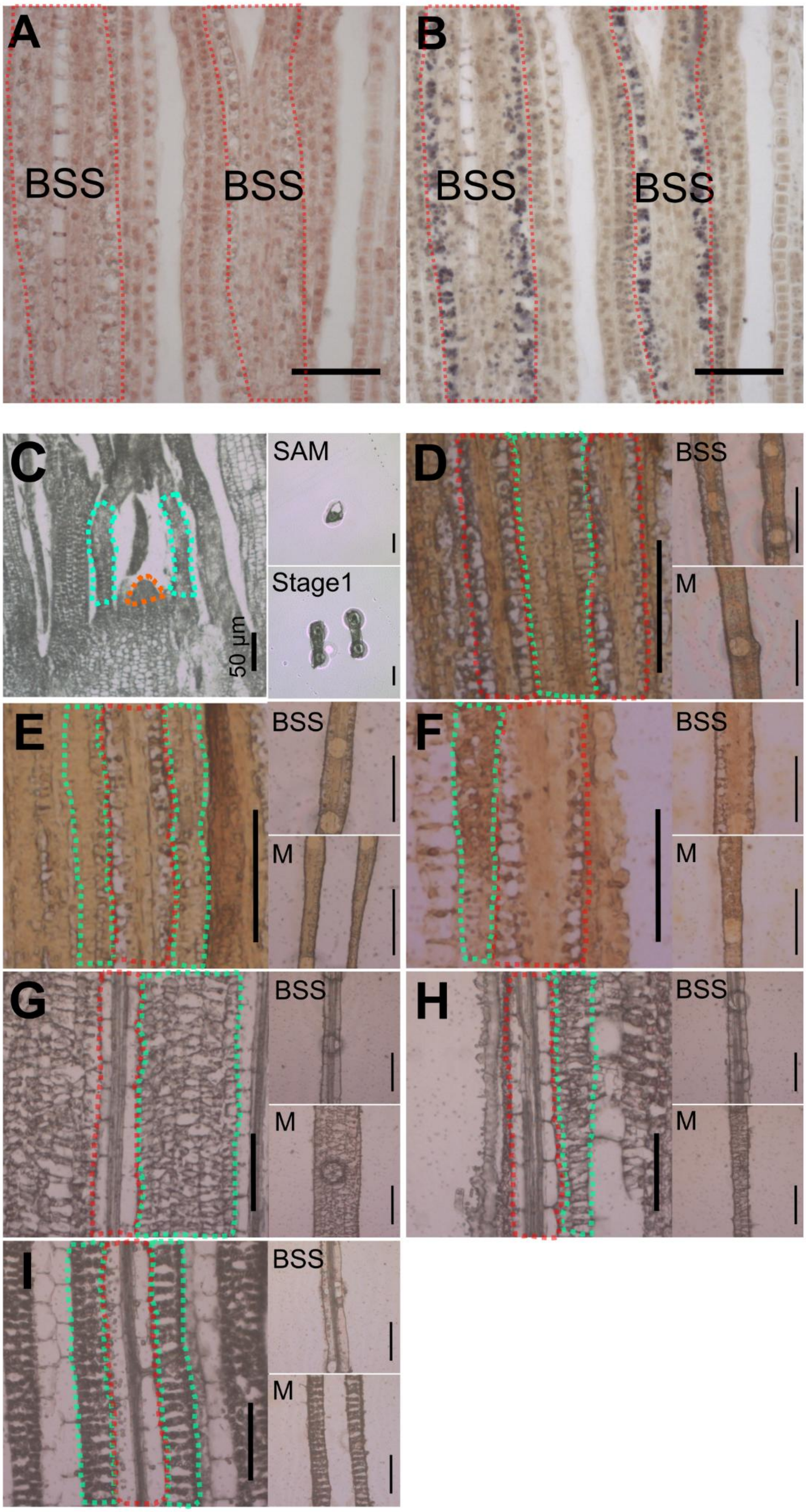

A
BSS
BSS
B
BSS
BSS
C
SAM
Stage1
50 μm
D
BSS
M
E
BSS
M
F
BSS
M
G
BSS
M
H
BSS
M
I
BSS
M

**Supplementary Figure 5. Starch staining allowed identification of bundle sheath cells prior to clear cell differentiation, and so allowed mesophyll and bundle sheath cells to be targeted for laser capture microdissection**.

(**A**,**B**) Paradermal sections at the base of rice young leaves (Stage 2) before iodine staining (**A**) in which bundle sheath and mesophyll cells are hard to discriminate. (**B**) After iodine staining bundle sheath chloroplasts were more darkly stained than mesophyll chloroplasts. M: mesophyll, BS: bundle sheath, V: veinal cells (including xylem and phloem cells). (**C-I**) Laser capture microdissection of Shoot Apical Meristem (SAM) and leaf primordia (Stage 1) (**C**), bundle sheath strands (BSS) and mesophyll (M) samples at stage 2 (**D**), 3 (**E**), 4 (**F**), 5 (**G**), 6 (**H**) and 7 (**I**) for transcriptomic analysis. Prior to microdissection, leaf sections from stage 2, 3, and 4 (**D-F**) were stained in 1% (w/v) iodine solution (in acetone) for 1 min. Scale bars: 50 µm. In **C**, SAM was marked with orange dash line, leaf four primordia in turquoise dash line, in **A**, **B** and **D-I**, bundle sheath cells were marked with red dash lines, mesophyll in green dash lines.

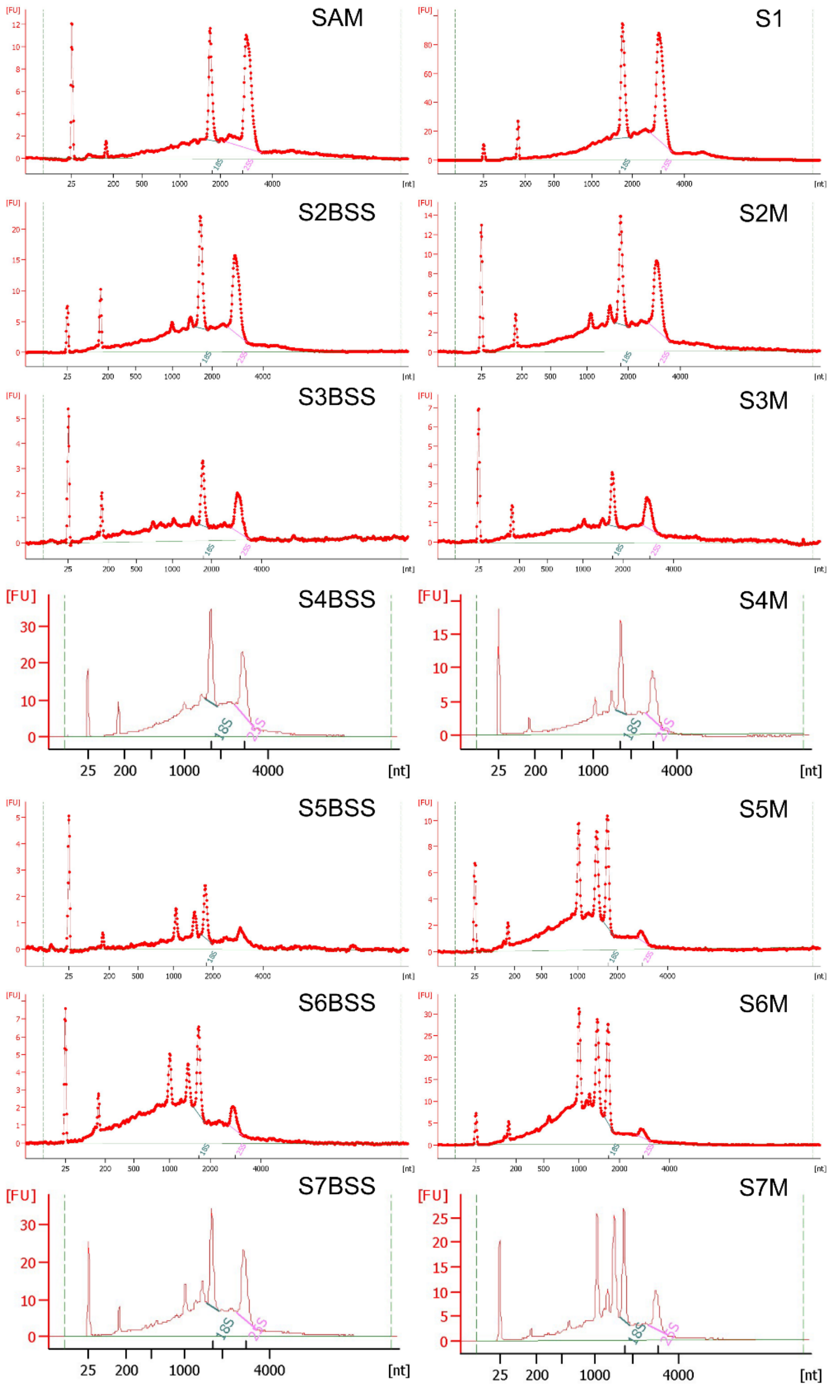


**Supplementary Figure 6. RNA isolated from shoot apical meristems, leaf primordia, bundle sheath strands and mesophyll cells across the developmental stages.**

Good RNA quality indicated by Bioanalyzer profiles, and while chloroplastic rRNA peaks were absent in samples from the shoot apical meristem (SAM) and leaf 4 primordia (S1), they were detectable in stages 2 to 4 (S2-S4) and subsequently clearly increased. This response was more pronounced in mesophyll samples.

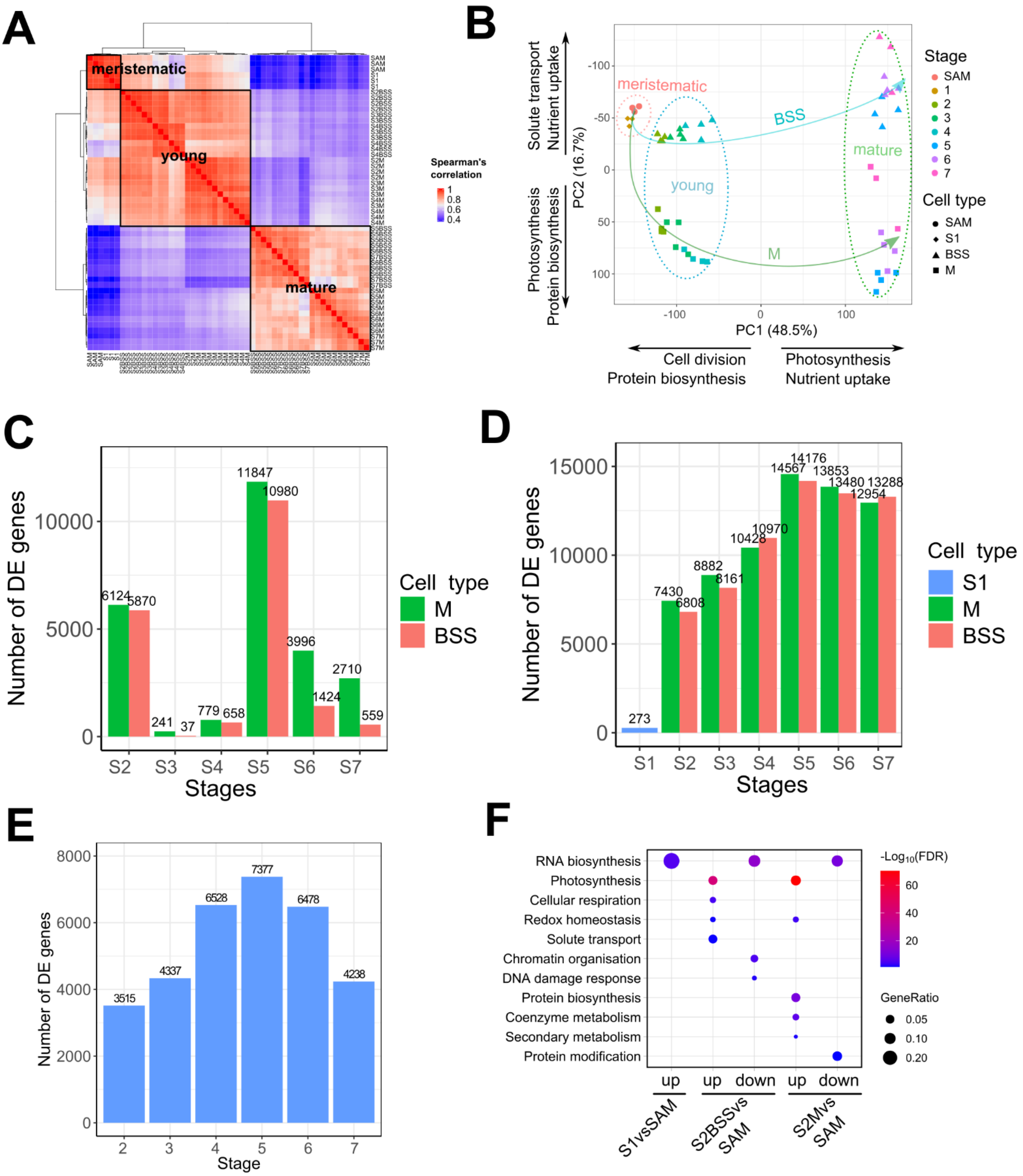
A
meristematic
young
mature
Spearman's correlation
1
0.8
0.6
0.4
B
Solute transport
Nutrient uptake
Photosynthesis
Protein biosynthesis
PC2 (16.7%)
-100
-50
0
50
100
meristematic
BSS
young
mature
M
-100
0
100
PC1 (48.5%)
Cell division
Protein biosynthesis
Photosynthesis
Nutrient uptake
Stage
SAM
1
2
3
4
5
6
7
Cell type
SAM
S1
BSS
M
C
Number of DE genes
10000
5000
0
6124
5870
241
37
779
658
11847
10980
3996
1424
2710
559
S2
S3
S4
S5
S6
S7
Stages
Cell type
M
BSS
D
Number of DE genes
15000
10000
5000
0
273
7430
6808
8882
8161
10428
10970
14567
14176
13853
13480
12954
13288
S1
S2
S3
S4
S5
S6
S7
Stages
Cell type
S1
M
BSS
E
Number of DE genes
8000
6000
4000
2000
0
3515
4337
6528
7377
6478
4238
2
3
4
5
6
7
Stage
F
RNA biosynthesis
Photosynthesis
Cellular respiration
Redox homeostasis
Solute transport
Chromatin organisation
DNA damage response
Protein biosynthesis
Coenzyme metabolism
Secondary metabolism
Protein modification
up
up
down
up
down
S1vsSAM
S2BSSvs SAM
S2Mvs SAM
-Log10(FDR)
60
40
20
GeneRatio
0.05
0.10
0.20

**Supplementary Figure 7. Global patterns gene expression in bundle sheath strands and mesophyll cells during leaf development.**

(**A**) Spearman's correlation of RNA-seq samples showed strong correlation between biological replicates and samples clustered into three main groups according to developmental stage, meristematic (SAM and stage 1), young (stage 2 to stage 4) and mature (stage 5 to stage 7). In young or mature leaf tissues samples were further grouped by cell type. (**B**) Principal component (PC) analysis indicated that the first two components of variance explained over 64% of variance, with the first capturing a strong developmental time component whereas the bundle sheath and mesophyll were clearly separated along the second axis. Bundle sheath and mesophyll samples displayed similar trajectories from the SAM and leaf primordia (S1), and a gap was evident between young and mature stages suggesting that significant remodelling had taken place at this time in both cell types. Functional enrichment of the top and bottom 5% loading genes showed that the first component was characterised by a reduction in transcripts associated with cell proliferation and protein synthesis but also increased photosynthesis and nutrient uptake activity as tissue matured, the second principal component was associated with solute transport and nutrient uptake for bundle sheath strands cells, but protein biosynthesis and photosynthesis in the mesophyll. (**C**) Number of differentially expressed genes in bundle sheath strands (BSS) and mesophyll (M) compared with each previous stage. (**D**) Number of differentially expressed genes of leaf primordia (S1), bundle sheath strands (BSS) and mesophyll (M) compared with SAM. (**E**) number of differentially expressed genes between BSS and M samples. (**F**) Functional enrichment analysis of differentially expressed genes between Shoot Apical Meristem (SAM) and leaf primordia (S1) or Stage 2 samples.

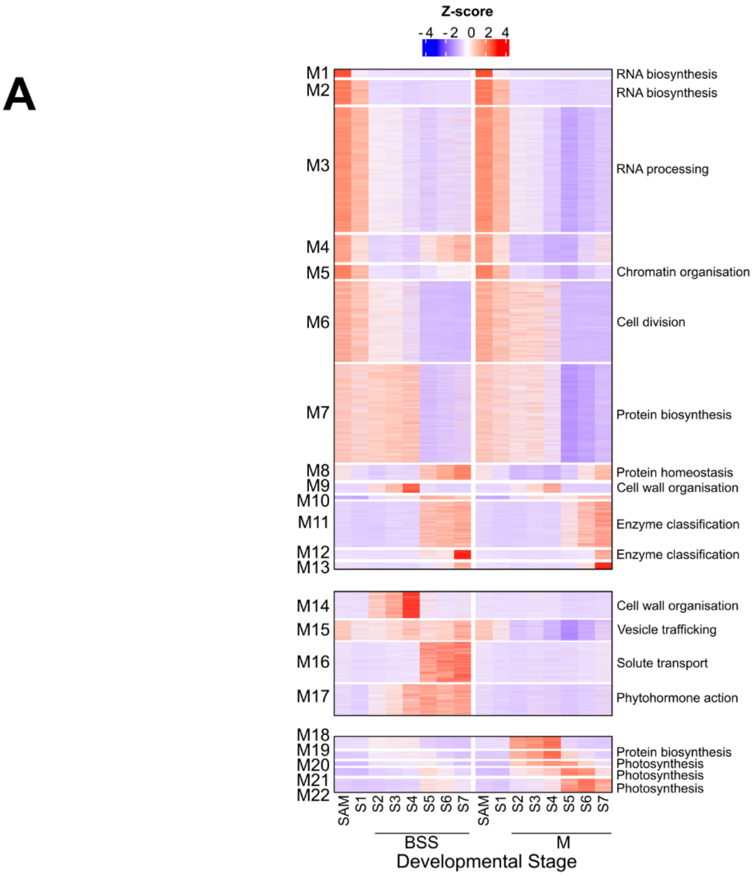


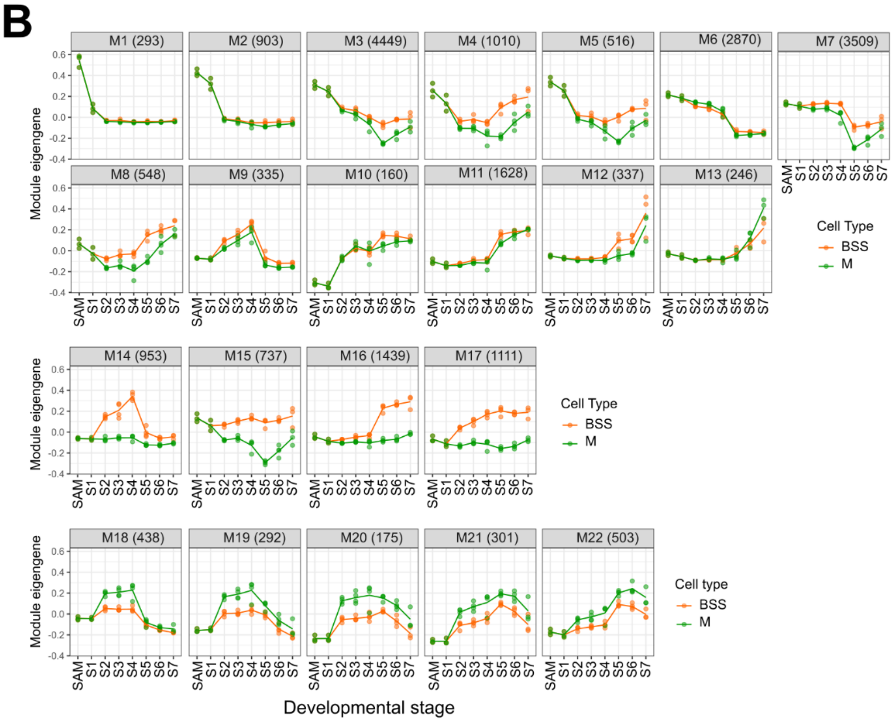


**Supplementary Figure 8. Expression patterns associated with the spatiotemporal sampling of mesophyll and bundle sheath cells in this study.**

(**A**) Heatmap depicting expression of 22,753 genes, and clustering identified three major patterns of gene expression. First modules with no obvious difference between bundle sheath and mesophyll (M1-13), second modules that were preferential to the bundle sheath (M14-17), and third modules preferential to mesophyll (M18-22) cells. The most enriched biological processes associated with each module are indicated on the right. Number of genes in each co-expression module indicated in parnetheses. (**B**) Expression patterns of module eigengenes which represents the first principal component of each co-expression module.

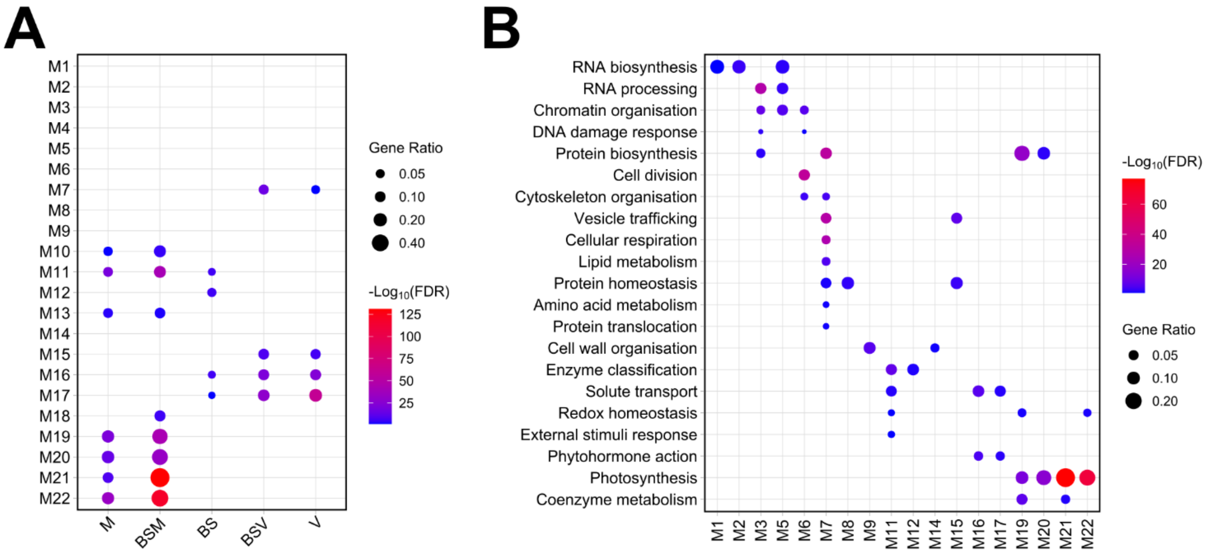


**Supplementary Figure 9. Cell type and functional enrichment of genes populating the 22 WGCNA modules.**

(**A**) over-representation analysis of genes associated with mesophyll (M), bundle sheath and mesophyll (BSM), bundle sheath (BS), bundle sheath and veins (BSV) and veins (V) (Hua et al., 2021). (**B**) Functional enrichment of Mercator 4 primary categories.

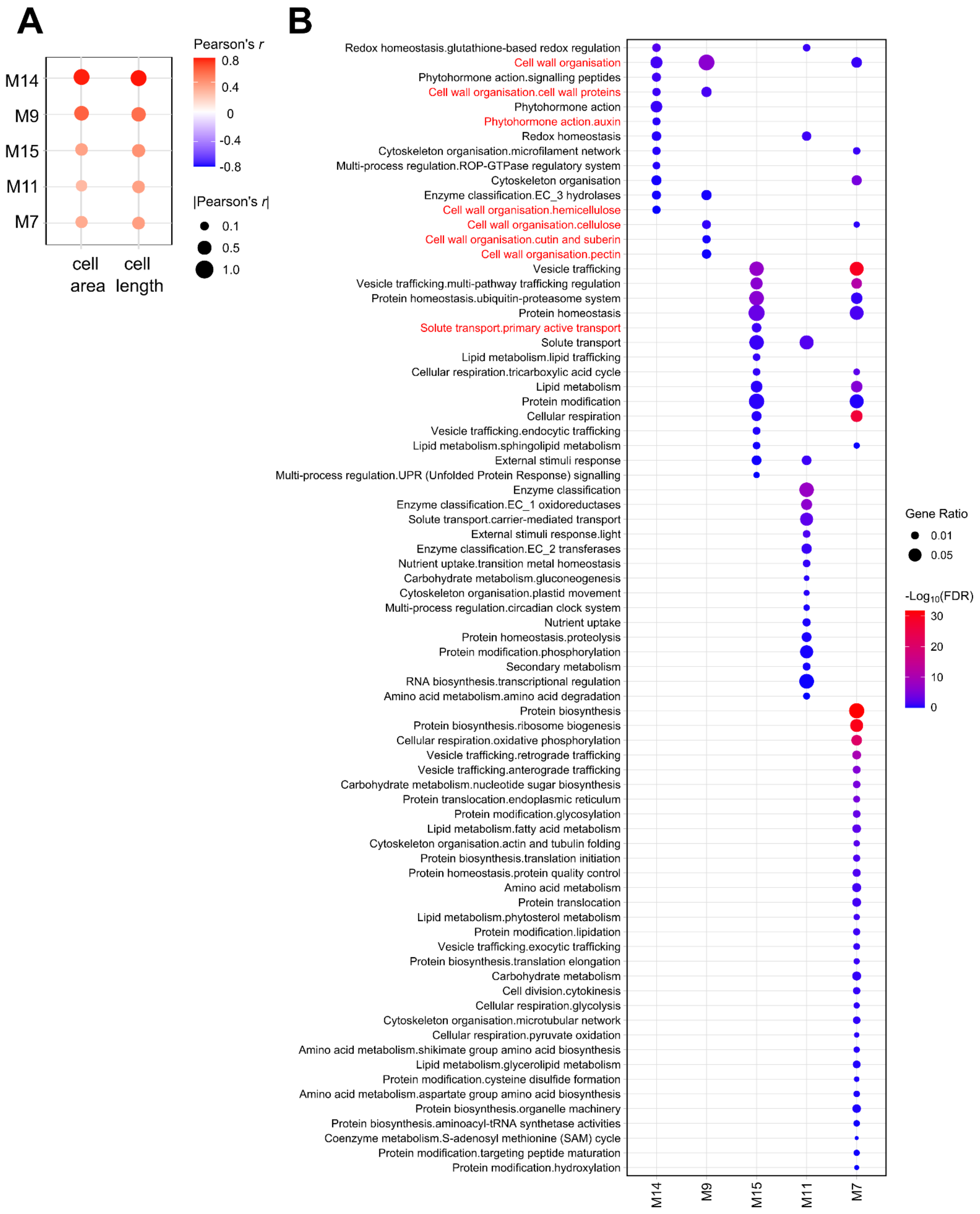


**Supplementary Figure 10. Transcripts associated with cell wall biosynthesis, auxin signalling were enriched in modules positively correlated with bundle sheath cell elongation.**

(**A**) Top five module eigengenes that were positively correlated with bundle sheath cell area and length. (**B**) Mercator 4 categories that were significantly over-represented in the five modules, categories associated with apoplastic acidification mediated cell expansion were highlighted in red.

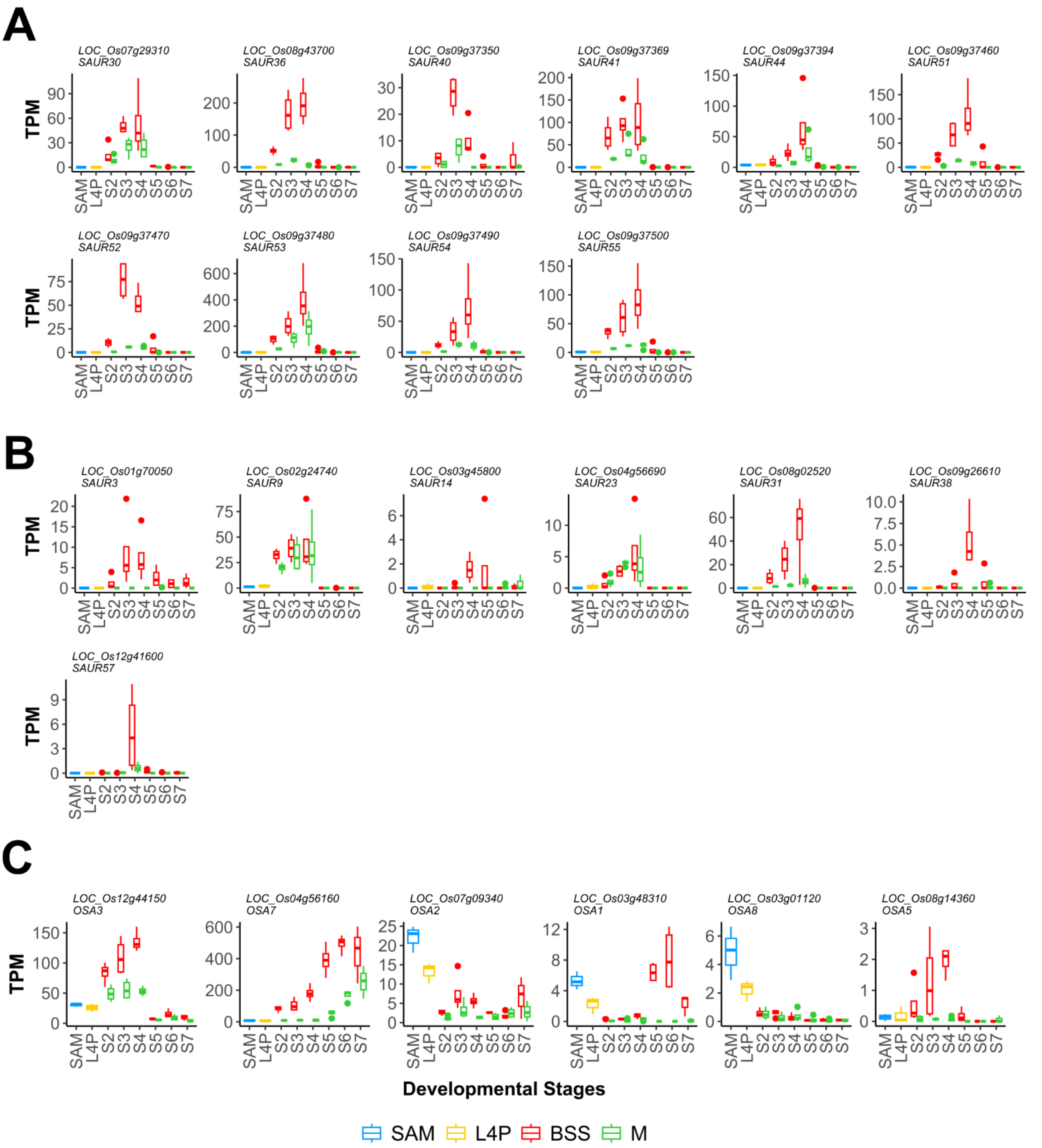


**Supplementary Figure 11. Transcript abundance for *SAUR* and plasma membrane *H⁺-ATPases* in bundle sheath strands (BSS) and mesophyll (M) cells.**

(**A,B**) *SAUR* genes from co-expression modules M14 and M9 - members closely related to Arabidopsis *SAUR63* shown in (**A**), and others in (**B**). (**C**) Six plasma membrane associated *H⁺-ATPase* genes detected in this dataset.

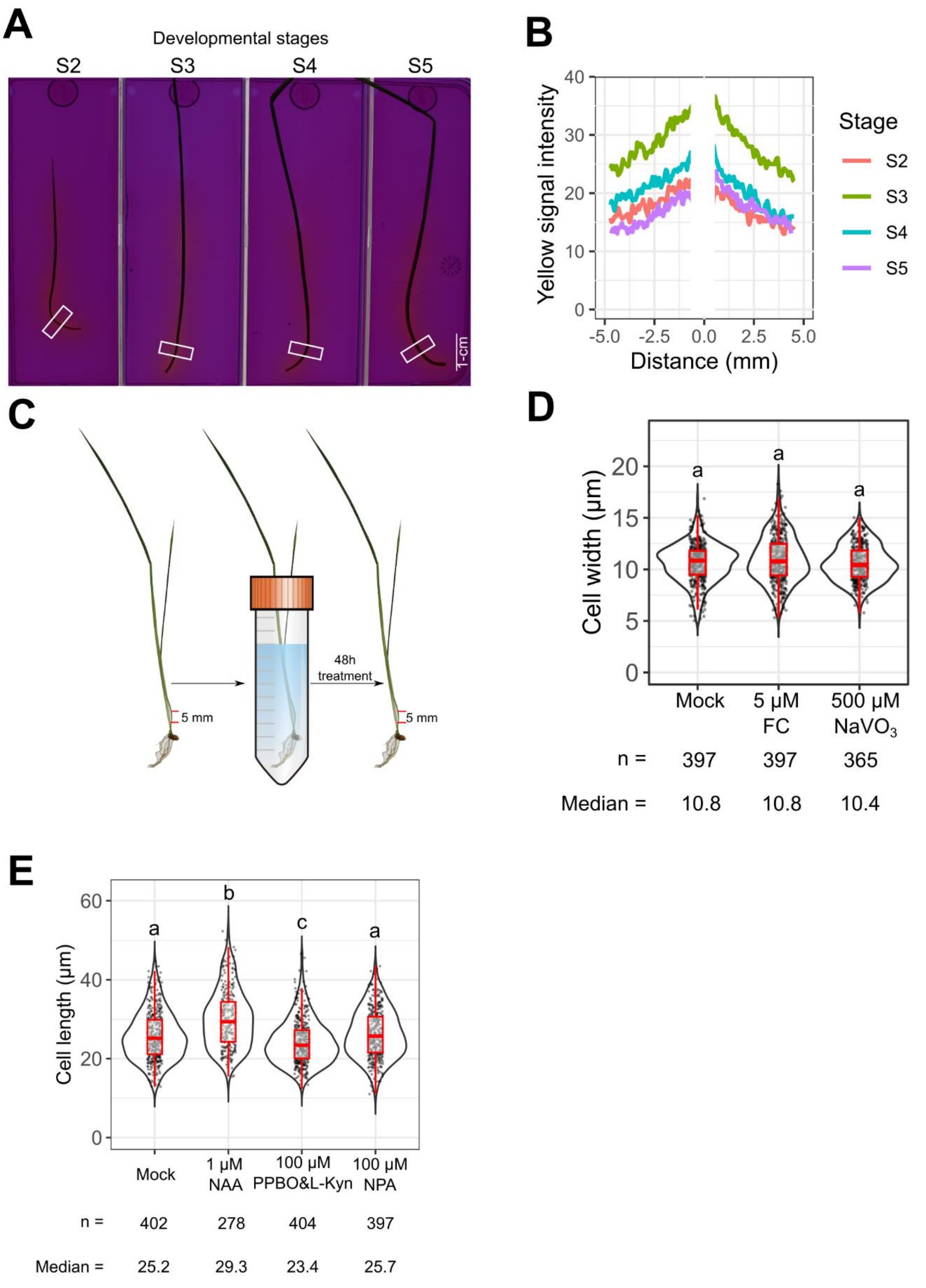


**Supplementary Figure 12. Summary of approach used and data derived after pharmacological interventions during bundle sheath development in rice leaves.**
(**A**) Basal regions of leaves at stage 2,3,4,5 are capable of acidifying agarose media (see gradient in colour as distance is increased from the leaf). (**B**) Quantification of acidification measured by grey value at the green channel (regions highlighted in A were measured). (**C**) Experimental set-up for pharmacological treatments. (**D**) Effect of fusicoccin and $NaVO_3$ on bundle sheath width. (**E**) Effect of auxin, inhibitors of auxin biosynthesis (PPBO & L-Kyn) or polar transport (NPA) on bundle sheath length. In **D**,**E**, data distribution was shown as violins plot, the 25$^{th}$, 50$^{th}$ and 75$^{th}$ percentile shown in boxplot with whiskers of 1.5x interquartile range. Different letters indicate statistically significant differences between treatments (FDR < 0.05) based on pairwise Wilcoxon rank-sum test with Benjamini-Hochberg correction.

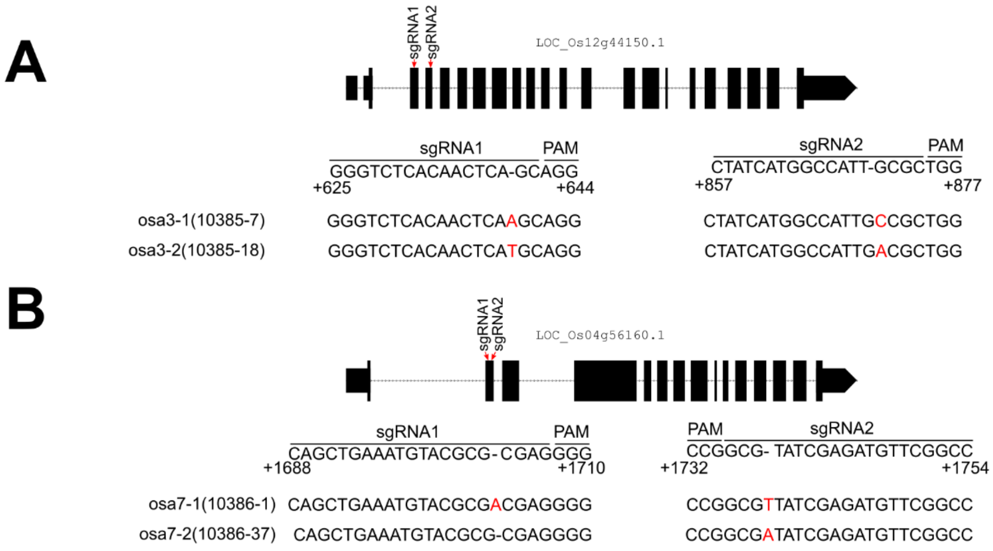


**Supplementary Figure 13. Design and genotyping after CRISPR editing of *OSA3* and *OSA7* mutants.**

(**A,B**) Two sgRNAs were designed to target the first or second exons of *OSA3* (**A**) or *OSA7* (**B**)*.* Homozygous mutations were identified close to the PAM sequences in each two mutant alleles for each gene by Sanger sequencing.

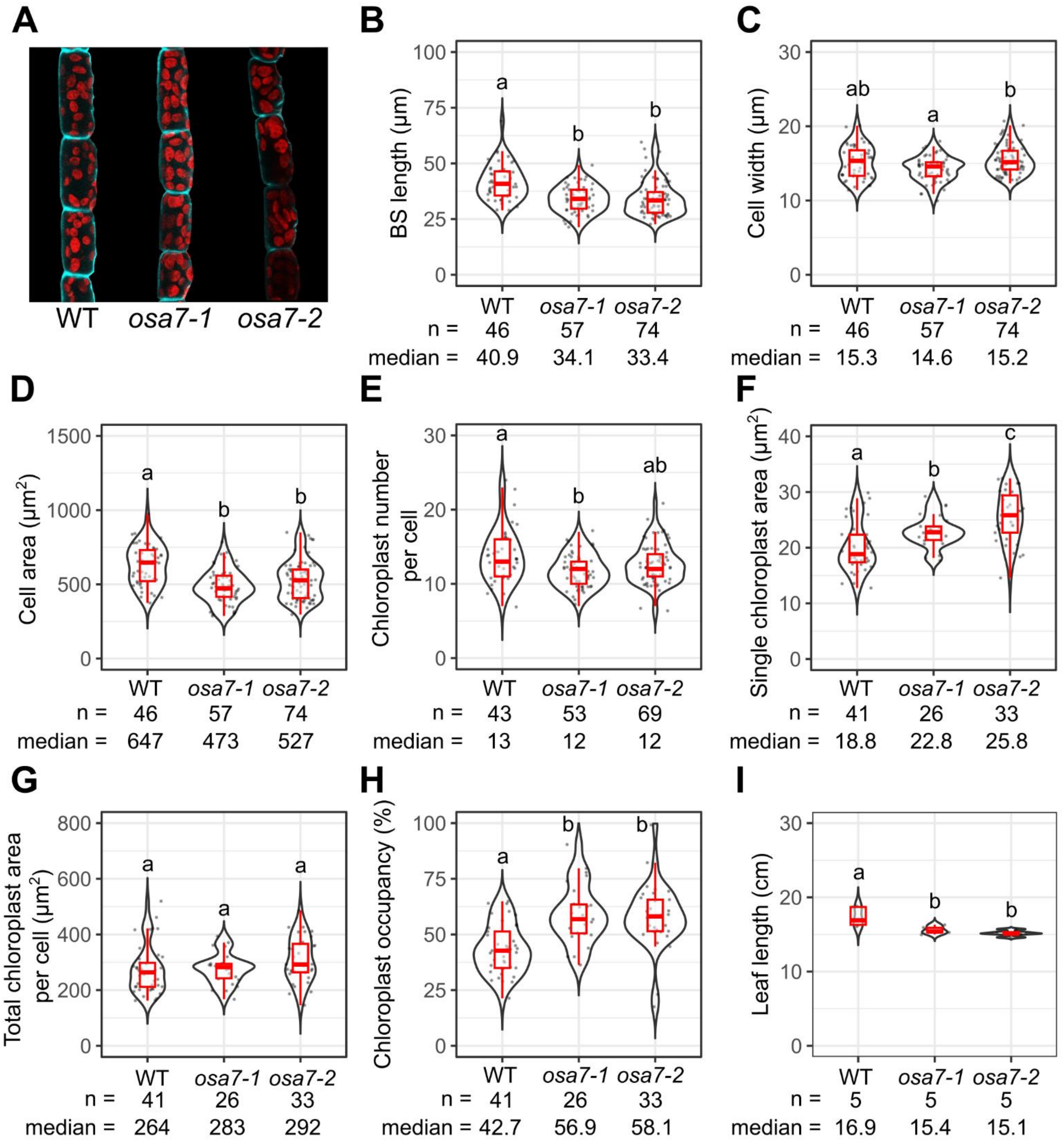


**Supplementary Figure 14. Quantification of cell and chloroplast parameters in *osa7* mutant alleles.**

(**A**) Representative images of bundle sheath cells: chlorophyll fluorescence indicated in red, cell wall stained by Calcofluor White and indicated in turquoise. (**B-I**) Impact of *OSA7* knock-out on bundle sheath length (**B**), width (**C**), bundle sheath cell area (**D**), chloroplast number in the bundle sheath (**E**), individual chloroplast area in bundle sheath cells (**F**), total chloroplast area per bundle sheath cell (**G**), chloroplast coverage in bundle sheath cells (**H**), and leaf four length (**I**).

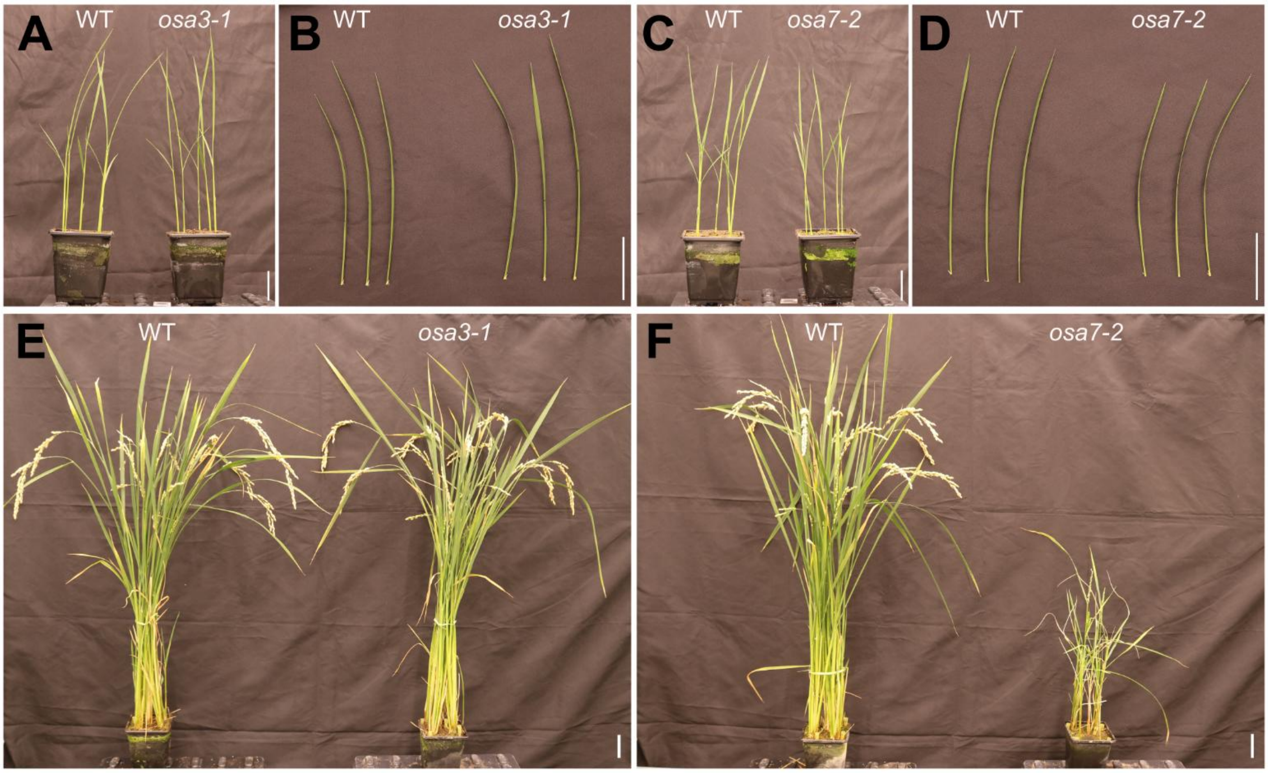


**Supplementary Figure 15. Growth of *osa3* and *osa7* mutants.**
(**A**, **C**) Representative images of 2-week-old *osa3* (**A**) and *osa7* (**C**) mutants. (**B**, **D**) Images of the fourth leaf from *osa3* (**B**) and *osa7* (**D**) plants. (**E**,**F**) 3-month-old *osa3* (**E**) and *osa7* (**F**) plants. Scale bars represent 5 cm.

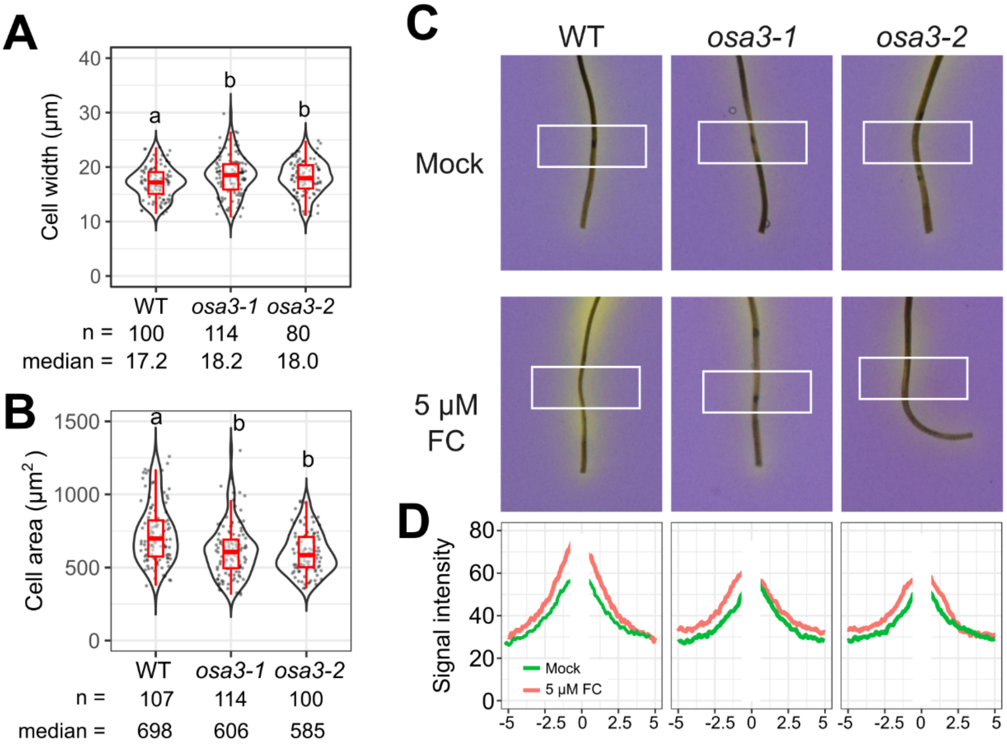


**Supplemental Figure 16.** ***osa3*** **mutants exhibit reduced sensitivity to fusicoccin.** (**A**,**B**) Quantification of bundle sheath width (**A**), bundle sheath cell area (**B**), (**C**) Acidification of agarose gel by stage 3 leaves from wild type and *osa3* mutants, visualized using bromocresol purple. (**D**) Quantification of acidification based on the extent of agarose gel yellowing within the white boxed region shown in (**C**).

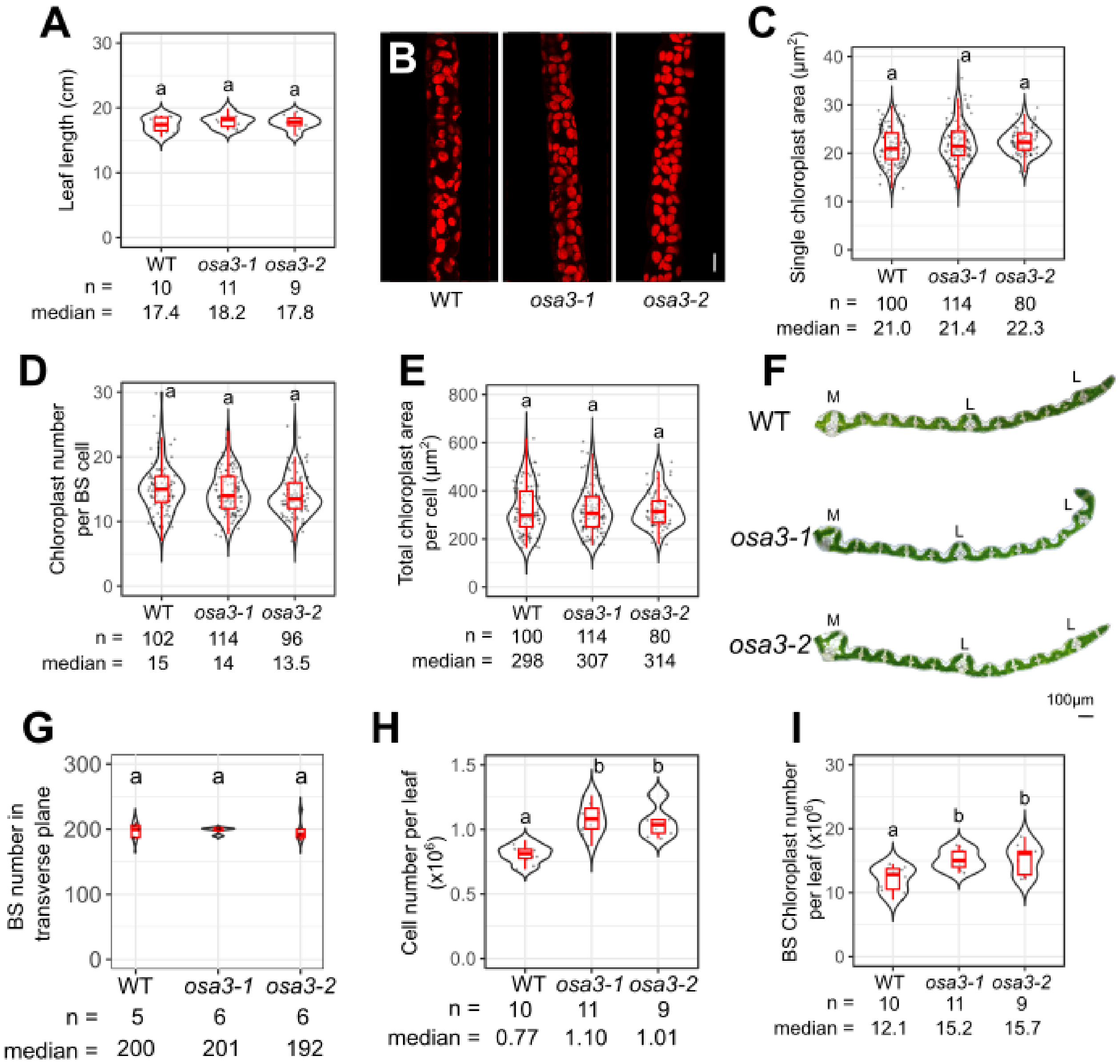


**Supplementary Figure 17. Impact of *OSA3* knock-out on number and chloroplast content of the bundle sheath.**

(**A**) leaf length showed no significant changes in *osa3* mutants compared with wild type (WT) plants. (**B**) bundle sheath chloroplasts indicated in chlorophyll fluorescence in red. (**C-E**) area of individual chloroplast from bundle sheath cells (**C**), chloroplast number per bundle sheath cell (**D**) and total chloroplast area per bundle sheath cell (**E**) showed no significant difference between *osa3* mutants and WT. (**F**) Cross sections of the fourth leaf showing number of veins. M, L indicate midrib and lateral veins respectively, Intermediate veins are not marked. Scale bar indicates 100 µm. (**G**) number of bundle sheath cells in transverse plane in *osa3* and WT plants. (**H, I**) *osa3* mutants were estimated to have significantly more bundle sheath cells (**H**) and total chloroplast number of bundle sheath cells (**I**) in leaf four compared with wild type plants.
